# Revisiting the role of cAMP in Drosophila aversive olfactory memory formation

**DOI:** 10.1101/2023.06.26.545795

**Authors:** Takashi Abe, Daisuke Yamazaki, Makoto Hiroi, Yutaro Ueoka, Yuko Maeyama, Tetsuya Tabata

## Abstract

In aversive olfactory conditioning in Drosophila melanogaster, an odor (the conditioned stimulus, CS) is paired with an electric shock (the unconditioned stimulus, US). Rutabaga adenylyl cyclase, which synthesizes cAMP in Kenyon cells (KCs), has long been proposed to act as a coincidence detector, synergistically activated by calcium/calmodulin in response to odor-evoked KC activity and by Gαs downstream of shock-evoked dopamine signaling. This framework implies that CS+ responsive KCs, which receive both CS and US signals, should be distinguishable from the remaining KCs by elevated cAMP levels, such that elevated cAMP would selectively mark these KCs for memory-related plasticity. However, our live imaging revealed that olfactory stimulation alone activates dopaminergic signaling and elevates cAMP broadly across KC axons, with only weak spatial correspondence to sparse local Ca^2+^ responses, such that CS+ responsive KCs could not be distinguished from the remaining KCs by bulk cAMP levels. This observation challenges a key prediction of the Rutabaga coincidence-detection model. Our findings instead suggest that cAMP concentrations do not selectively mark CS+ neurons. Rather, we propose that the compartmental population dynamics of cAMP across the γ lobe represent a KC-intrinsic expression of valence, integrated across compartments from multiple dopaminergic inputs. Associative conditioning reshaped dopamine responses to the CS+ odor, with corresponding changes in this compartmental cAMP pattern. Simultaneous imaging of cAMP and acetylcholine (ACh) revealed that conditioning establishes an inverse relationship between this pattern and ACh release for the CS+ but not for the CS− odors; this relationship was absent in rutabaga mutants. Pharmacological elevation of cAMP suppressed ACh release from KCs. These findings suggest that cAMP-dependent suppression of ACh release from KCs may shift the valence of the CS+ odor toward a more aversive state in conditioned flies, thereby contributing to aversive olfactory memory formation.

## Introduction

A quintessential aspect of classical conditioning involves the temporal pairing of an initially neutral stimulus, the conditioned stimulus (CS), with a reinforcing stimulus, the unconditioned stimulus (US), and the preservation of this association at the level of synaptic connections, leading to the manifestation of a conditioned response to the CS (Rescorla, 1988). The temporal correlation between the CS and US is mediated by coincidence detection mechanisms. One well-known coincidence detector is adenylyl cyclase (AC), which is stimulated by both calcium/calmodulin and receptor-coupled G-proteins and catalyzes the conversion of ATP to cAMP. This role was first revealed by the pioneering studies on classical conditioning of gill-withdrawal reflex in the marine mollusk *Aplysia*. When siphon stimulation (CS) and an electric shock to the tail (US) were repeatedly paired, the CS elicited potentiated gill- and siphon-withdrawal reflexes (Carew et al., 1981). Integration of the calcium influx resulting from the CS and the serotonin release resulting from the shock to the tail (US) increased cAMP levels (Ocorr et al., 1985), as also shown biochemically (Abrams and Kandel, 1988; Yovell et al., 1992). These physiological and biochemical studies highlighted the pivotal role of adenylyl cyclase as a coincidence detector. This view was further strengthened by the genetic dissection of Drosophila olfactory conditioning (Abrams and Kandel, 1988; Levin et al., 1992; Livingstone et al., 1984; Tully, 1987; Tully and Quinn, 1985). Odor information via acetylcholine (ACh) signaling represents the CS (Gu and O’Dowd, 2006), and G-proteins coupled to dopamine metabotropic receptors (Kim et al., 2007) activated by dopaminergic neurons encode the US (electric shock or sugar reward) in Drosophila (Burke et al., 2012; Liu et al., 2012; Riemensperger et al., 2005; Schwaerzel et al., 2003). These two input pathways converge on the Rutabaga adenylyl cyclase (AC) (Davis, 1993; Livingstone et al., 1984) expressed in Kenyon cells (KCs) (McGuire et al., 2003; Zars et al., 2000) of the mushroom bodies (MBs) (Heisenberg, 2003).

Genetically encoded fluorescent sensors have recently enabled the microscopic visualization of the dynamics of AC activity in fly brains. Tomchik and Davis showed the Rutabaga AC-dependent synergistic increase in cAMP in α/β and α′/β′ KCs when acetylcholine (CS) and dopamine (US) application were paired to ex vivo brain explants (Tomchik and Davis, 2009). They also demonstrated that elevating cAMP acutely enhanced KC excitability. Additionally, Handler et al. reported a synergistic increase in cAMP levels in γ KCs upon temporal pairing of acetylcholine application with dopamine neuron activation in ex vivo brain explants (Handler et al., 2019). Gervasi et al. employed protein kinase A (PKA) activity as an indicator of cAMP levels, and reported that the simultaneous administration of acetylcholine and dopamine led to an augmentation of PKA activity exclusively in α/β KCs in a manner dependent on the Rutabaga AC (Gervasi et al., 2010). These findings collectively indicate that the Rutabaga AC fulfills its long-held role as a coincidence detector. It is worth noting, however, that these imaging experiments did not measure cAMP or PKA transients at cellular resolution, but rather as a unit of the lobe, comprising a cluster of KC axons. It is estimated that approximately 5–9% of KCs exhibit a consistent response to monomolecular odors such as 3-octanol (OCT) and 4-methylcyclohexanol (MCH), as employed in these experiments (Honegger et al., 2011). This sparse nature of coding is essential for distinguishing the odor paired with the US (CS+) from an unpaired odor (CS−) in classical conditioning. Thus, if the Rutabaga AC serves as the coincidence detector, cAMP levels in odor-responsive KCs activated by the CS+ should be preferentially elevated upon pairing with dopamine input from the US, relative to those in the remaining KCs. Recent studies demonstrated that the axo-axonic connections mediated by the muscarinic type-B receptor exert a suppressive effect on dopamine-evoked cAMP in neighboring KCs, thereby preventing weakly activated and unreliably responding KCs (off-target KCs) from being erroneously classified as the genuine CS+ responsive KCs. This phenomenon of lateral inhibition is deemed essential for memory formation (Manoim et al., 2022; Wolkovitz et al., 2026). To the best of our knowledge, there is no direct evidence that the cAMP levels in CS+ responsive KCs are higher than those in the remaining KCs following classical conditioning. However, such selective elevation would be a crucial indicator that AC functions as the coincidence detector. Therefore, live imaging should ideally be conducted at a cellular resolution, with the odors and electric shocks delivered as the CS and US, respectively, to preserve the sparse nature of coding and the physiological authenticity of the signals. With this objective in mind, we established an imaging system that simultaneously records cAMP and calcium transients in the KCs of head-opened Drosophila melanogaster under a two-photon microscope. We verified that odor presentation (CS) evoked not only calcium influx but also dopamine release (Mao and Davis, 2009; Riemensperger et al., 2005) and discovered that it induced robust cAMP increases even in the absence of the US, including in axons that did not exhibit elevated calcium levels. This substantial cAMP elevation, which was dependent on dopamine signaling but was largely uncoupled from local calcium concentrations, raised questions about the long-standing model of coincidence detection in olfactory conditioning. Based on reevaluation of CS- and US-evoked cAMP, calcium, and dopamine transients, downstream PKA activity, and acetylcholine (ACh) release from KCs, we discuss what has been overlooked in the Drosophila model of classical conditioning. We further propose that compartmental patterns of cAMP across the γ lobe represent a KC-intrinsic expression of integrated valence, which is reorganized for the CS+ by aversive conditioning and contributes to the suppression of ACh release from KCs.

## Results

### Odor presentation evokes broad, compartmental cAMP elevation across KCs, largely uncoupled from sparse Ca^2+^ responses

The mushroom bodies (MBs) in the Drosophila brain are the locus of olfactory classical conditioning (Heisenberg, 2003). The ∼2,000 KCs of the MBs are classified into three subtypes that extend axons into the α/β, α′/β′, and γ lobes (Crittenden et al., 1998)(Figure 1A). These lobes can be anatomically and functionally subdivided into 15 non-overlapping compartments, each innervated by distinct dopamine neuron (DAN) populations (Aso et al., 2014; Tanaka et al., 2008). In Drosophila olfactory conditioning, Rutabaga adenylyl cyclase (Rut AC) is thought to serve as the coincidence detector in KCs (McGuire et al., 2003; Zars et al., 2000)(Figure 1B). Dubnau and his colleagues have shown that Rutabaga adenylyl cyclase expression is required in both α/β KCs and γ KCs to support aversive memory fully (Blum et al., 2009), while the dopamine receptor Dop1R1 expression in γ KCs is sufficient to support aversive memory (Qin et al., 2012). In addition, protocerebral posterior lateral 1 (PPL1) cluster DANs projecting to the γ1, γ2, and α3 compartments and protocerebral anterior medial (PAM) cluster DANs projecting to the γ3 compartment of KCs convey aversive reinforcement signals (Aso et al., 2012; Yamagata et al., 2016). Elevations of cAMP levels and protein kinase A (PKA) activity reportedly exhibited a synergistic effect following the pairing of a conditioned stimulus with an unconditioned stimulus in the α lobe in the context of aversive conditioning paradigms (Gervasi et al., 2010; Tomchik and Davis, 2009), and in the γ lobe in the context of appetitive conditioning paradigms (Handler et al., 2019). Thus, we focused on the γ and α lobes in subsequent live imaging experiments. Calcium transients and cAMP levels evoked by odors were simultaneously measured using the genetically encoded fluorescent sensors jRGECO1b (Dana et al., 2016) and G-Flamp1 (Wang et al., 2022), respectively.

**Figure 1.**
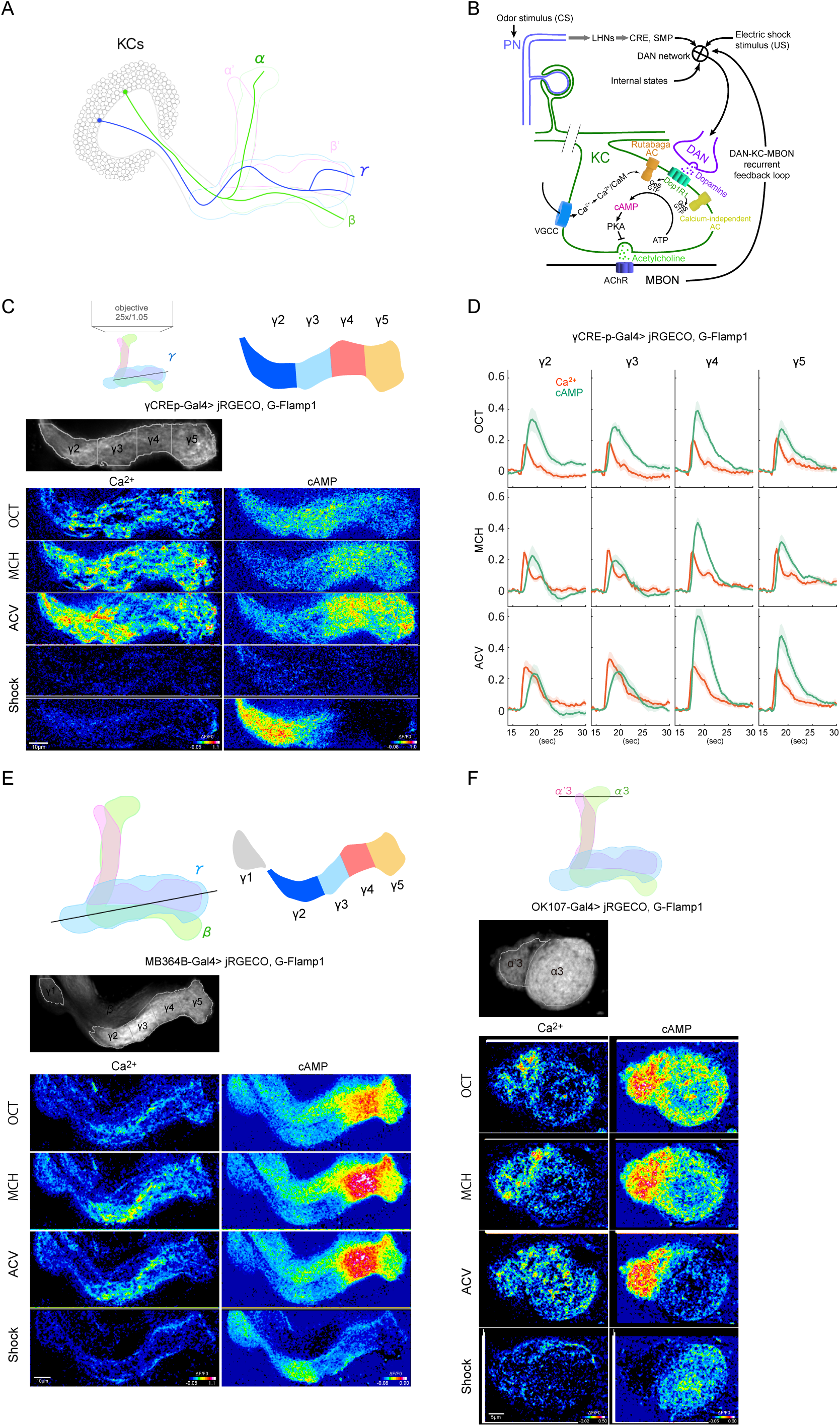
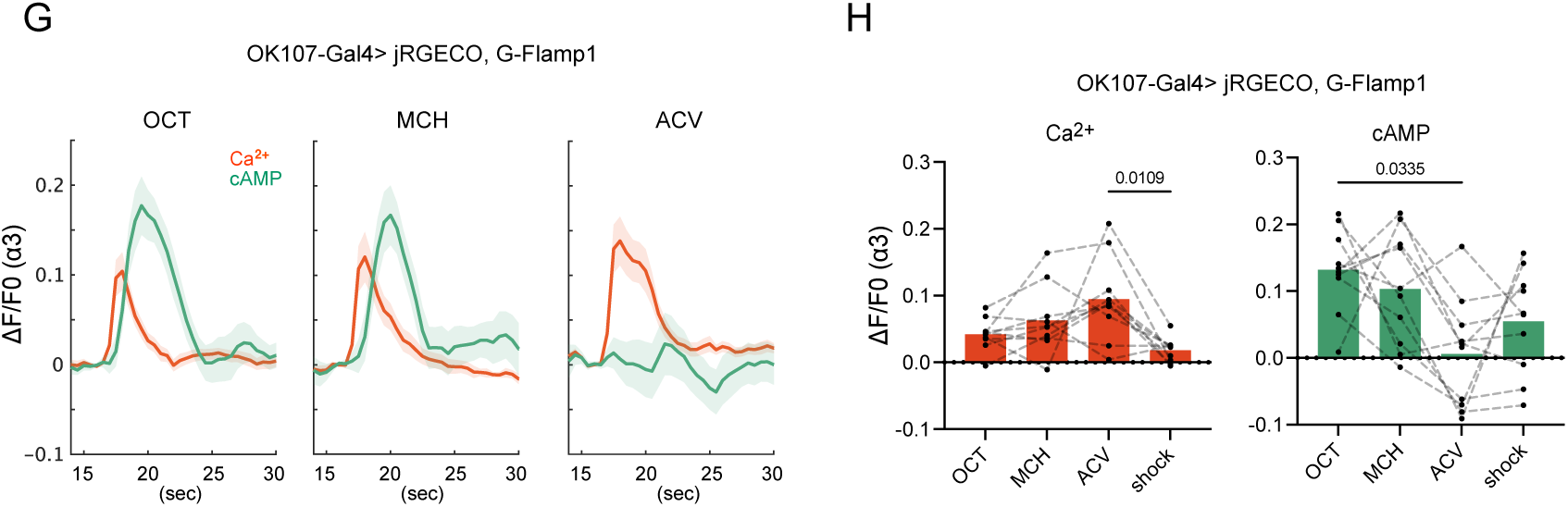
Odor presentation evokes broad, compartmental cAMP elevation across KCs, largely uncoupled from sparse Ca^2+^ responses. (A) Schematic of Kenyon cells (KCs), adapted from (Bilz et al., 2020). The cell body, the α/β, α′/β′ and γ lobes, and the α/β and γ KCs are depicted. (B) Schematic of the proposed role of Rutabaga adenylyl cyclase (Rut AC) as a coincidence detector. Odor presentation as the conditioned stimulus (CS) activates acetylcholine signaling in KCs and elevates intracellular Ca^2+^ through voltage-gated calcium channels. Aversive stimuli, such as electric shock, activate PPL1 DANs, which provide dopaminergic input to KC axons in the MB lobes. PPL1 DAN activity is shaped by odor information and noxious stimuli through brain regions including the lateral horn neurons (LHNs), crepine (CRE), and superior medial protocerebrum (SMP) (Kato et al., 2023), and is also regulated by the DAN-KC-MBON recurrent feedback loop (Adel and Griffith, 2021). The CS- and US-associated pathways can converge on Rut AC, which is synergistically activated by the calcium/calmodulin complex and the Gαs subunit of the G-protein complex coupled to the D1-like dopamine receptor Dop1R1. (C) Schematic representation of in vivo imaging of KCs. The objective and frontal view of the KCs along with the focal plane (upper left) and the dorsal view of γ KCs with color-coded γ2-5 compartments (upper right) are shown. Two-photon live imaging of the γ2-γ5 compartments of KCs was performed in flies expressing jRGECO1b and G-Flamp1 under the control of the γCRE-p-GAL4 driver. Exemplary, finely resolved images are shown. Averaged calcium responses during a 40-second air exposure were used to show the regions of interest (ROIs) for each compartment, depicted in grayscale. Pseudocolor images show averaged peak calcium and cAMP responses (10 frames for 5 s) to 3-octanol (OCT), 4-methylcyclohexanol (MCH), apple cider vinegar (ACV), and electric shock (shock). Two separate single-shock trials are shown. All images in the present study were acquired using a dual-channel line-scanning mode. (D) The graph shows the changes in calcium (in red) and cAMP (in green) levels over 15–30 s in response to different odors (OCT, MCH, and ACV) in four γ lobe compartments (γ2-γ5) driven by the γCRE-p-GAL4 driver. The odors were presented by opening the solenoid valve between 15-18 seconds. Solid lines represent the average response from 8 flies, and shaded areas show the standard error of the mean (SEM). (E) The focal plane of live imaging for compartments γ1-γ5 is schematically presented (Upper panel). Two-photon live imaging on the γ1-γ5 compartments of KCs in flies expressing jRGECO1b and G-Flamp1, driven by the MB364B-GAL4 driver. Averaged calcium responses during a 40-second air exposure were utilized to show ROIs for each compartment, depicted in grayscale. The ΔF/F0 images of averaged peak responses (10 frames, 5 s) of calcium and cAMP during the presentations of OCT, MCH, ACV, and shock in compartments γ1-γ5 (lower panels). (F) The focal plane of live imaging for the α3/α′3 compartments is schematically presented (upper panel). Two-photon live imaging on the α3/α′3 compartments of KCs in flies expressing jRGECO1b and G-Flamp1, driven by the OK107 driver. Averaged calcium responses during a 40-second air exposure were utilized to show ROIs for each compartment, depicted in grayscale. Magnified images of averaged peak ΔF/F0 responses (10 frames, 5 s) of calcium and cAMP in the α3 and α′3 compartments are shown. (G) The graph shows the changes in calcium (in red) and cAMP (in green) levels over 15–30 s in response to different odors (OCT, MCH, and ACV) in the α3 lobe compartment driven by the OK107-GAL4 driver. The odors were presented by opening the solenoid valve between 15-18 seconds. Solid lines represent the average response from 10 flies, and shaded areas show the standard error of the mean (SEM). ACV elicited the highest calcium response but the lowest cAMP response, compared to OCT and MCH. (H) The ΔF/F0 images, obtained by averaging the responses to odor stimuli over a duration of 5 seconds (10 frames), were quantified. The plots of the α3 compartment derived from the same sample are connected with dashed lines. Bar graphs represent mean values. Statistical comparisons across the four stimulus conditions (OCT, MCH, ACV, and shock) were performed using a Friedman test followed by Dunn’s multiple-comparisons test. p-values are shown only for significant pairwise comparisons among the four stimulus conditions (p < 0.05; n = 10). Figure 1C-video (related to Figure 1C and D). Figure 1D-video (related to Figure 1D). Figure 1E-video (related to Figure 1E). Figure 1F-video (related to Figure 1F, G and H). Figure 1G-video (related to Figure 1G and H). Videos illustrating simultaneous calcium and cAMP imaging, acquired at 2 Hz and played back at 6 Hz. The odor was presented for 3 seconds, from 16 to 19 seconds after the onset of imaging (corresponding to frames 32–38), and the electric shock was delivered at 17.5 seconds (frame 35).

The two sensors were initially expressed with the γCRE-p GAL4 driver (Yamazaki et al., 2018), as it labels a smaller subset of γ KCs compared to other γ KC drivers, enabling near–single-cell resolution within a dense neuropil. Specifically, the γCRE-p driver covers approximately 350 (Yamazaki et al., 2018) out of 701 γ KCs (Li et al., 2020). The γ lobe comprises five compartments, with the γ1 compartment being the most lateral. Initially, imaging of the γ2-5 compartments was performed to achieve the optimal balance of resolution and scanning speed (Figure 1C). In general, a sequence of images, with a consistent focus, was captured at a frequency of approximately 2 frames per second. Three olfactory stimuli—3-octanol (OCT), 4-methylcyclohexanol (MCH), and apple cider vinegar (ACV)—were each presented for 3 seconds, and a 10-ms electric shock was also delivered. Prominent calcium responses were observed in a limited number of axons (Figure 1C), reflecting the sparse coding nature of KCs (Honegger et al., 2011). The γ KC axons branch and traverse the lobe in a meandering manner, as depicted schematically in Figure 1A, resulting in single-plane images of calcium responses often appearing interrupted and patchy. As previously reported (Wang et al., 2022), we also detected odor-evoked cAMP signals with the second channel. These signals varied in a compartment-specific manner along the length of the lobes with distinct characteristics for each odor, and the most robust signal was identified within the γ4 compartment upon MCH or ACV presentation (Figure 1D). Notably, there was no evidence that cAMP was selectively synthesized in odor-responsive (as marked by the calcium signals in Figure 1C) axons. A marked increase in cAMP was temporally delayed relative to the rise in Ca^2+^ levels in a population of axons spanning almost the entire width of compartments, although there were variations in levels among individual compartments. Conversely, the calcium signal was sparsely distributed, indicative of a restricted number of KCs responding to a specific odor. These profiles suggest that cAMP levels do not exhibit a proportional relationship with calcium concentrations, but rather are regulated as a compartmental unit. An electrical stimulus elicited cAMP synthesis in the γ2 compartment, albeit with variable signal strength (Figure 1C). Thus, we carefully monitored each fly during electrical stimulation to verify a behavioral response to the shock. Note that only a minimal increase in calcium levels was detected following electrical stimulation. Additionally, the split GAL4 driver MB364B, which is expressed in α/β and γ KCs, was used to eliminate any potential bias in cAMP signaling, as the γCRE-p driver contains 12 tandem repeats of the cAMP-responsive element (CRE) in its regulatory region (Yamazaki et al., 2018) and may be more likely to drive a population of cells responsive to cAMP signaling. The γ1-γ5 compartments were similarly subjected to imaging upon exposure to comparable olfactory and electrical stimuli. The calcium and cAMP transient profiles in MB364B-positive γ KCs were similar to those observed in γCRE-p KCs (Figure 1E). A marked increase in cAMP levels was noted in axons traversing nearly the full breadth of the γ lobe, whereas the calcium signal was only discernible in a limited number of axons. An electrical stimulus elicited cAMP synthesis in the γ1 and γ2 compartments, and in γ3 to a lesser extent. Notably, ACV evoked a relatively γ4-dominant cAMP pattern, whereas electric shock predominantly elevated cAMP in the γ1–γ3 compartments. Responses in β KCs were also imaged using the MB364B driver; however, they were not further analyzed. We subsequently used the OK107 driver to visualize the α3 compartment, which was similarly subjected to olfactory stimuli and electrical stimulation. A transverse optical section of the densely packed neuropil in the α3 compartment enabled the observation of all α KCs within a single focal plane. In this view, cAMP levels increased considerably in the majority, if not all, of KCs upon exposure to MCH or OCT, whereas uniformly minimal responses were detected upon exposure to ACV, regardless of individual KCs’ odor-specific calcium responses (Figure 1F-H), consistent with compartmental regulation of cAMP synthesis. In summary, the cAMP levels display odor-specific and compartment-dependent patterns, regardless of calcium levels. We therefore focused our subsequent analyses on γ KCs, as the γ1 and γ2 compartments are the sites of short- and mid-term aversive memory formation (Aso et al., 2010; Aso and Rubin, 2016).

### Stimulus-evoked dopamine signaling elevates cAMP in KCs

We examined whether the dopamine signal contributes to the cAMP synthesis elicited by odor stimulation, given that odor-evoked cAMP transients vary among odors and odors activate DANs (Figure 1B) (Cervantes-Sandoval et al., 2017; Mao and Davis, 2009; Riemensperger et al., 2005; Sun et al., 2020). The broad elevation of cAMP across KCs within each compartment may reflect dopamine release through volume transmission (Liu et al., 2021). We subsequently employed GRAB_DA2m (Sun et al., 2020) and Pink Flamindo (Harada et al., 2017) simultaneously in the γ lobe of γCRE-p-GAL4-labeled KCs to assess the spatial correspondence between the dopamine and cAMP signals, respectively. An example recording showed that dopamine and cAMP signals were both broadly distributed within the compartments (Figure 2A and D), in contrast to the sparse local calcium responses observed in KCs (Figure 1C, E, and F). The cAMP signal peaked later than the dopamine signal (Figure 2G). To further elucidate the role of dopamine, we measured the cAMP levels in a functionally null mutant of the type I dopamine receptor, Dop1R1 (Akiba et al., 2020), which couples to Gαs (Sugamori et al., 1995) and thus engages adenylyl cyclase activation. The cAMP signals were substantially attenuated in the Dop1R1 mutant, suggesting that dopamine signaling plays a predominant role in this activation (Figure 2B, C, E and F). In some recordings from Dop1R1^KO^ mutant flies, the shock-evoked cAMP response in the γ1 compartment, when detectable, appeared delayed relative to controls (Figure 2C). It is worth noting that the simultaneously recorded calcium signal was significantly augmented when odors were presented, in comparison to the genetic control (Figure 2H). These experiments suggest that odor stimulation alone evokes cAMP synthesis via dopamine released from both PPL1 DANs (Figure 1B), which project to the γ1 and γ2 compartments, and PAM DANs, which project to the γ3-γ5 compartments. Given that PPL1 DANs mediate reinforcement by electric shock, whereas many PAM DANs convey reward-related reinforcement, odor presentation as the CS may already recruit neural pathways involved in US processing.

**Figure 2.**
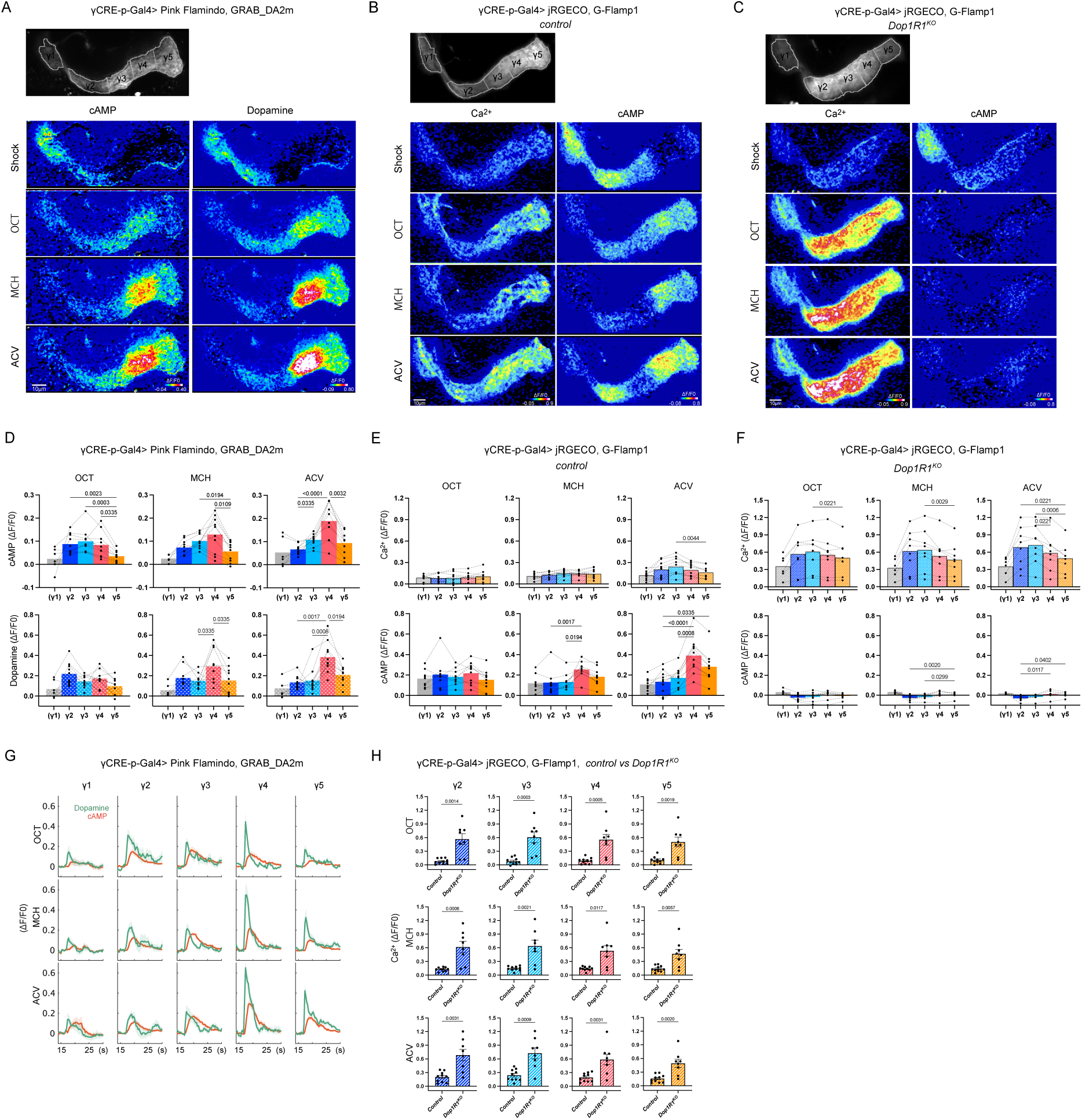
Stimulus-evoked dopamine signaling elevates cAMP in KCs. (A) Two-photon live imaging on γ1-γ5 compartments of KCs in flies expressing Pink Flamindo and GRAB_DA2m, driven by the γCRE-p-GAL4 driver. An averaged raw fluorescence image acquired during a 40-s air exposure was used to delineate the ROIs for each compartment and is shown in grayscale. ΔF/F0 images of averaged peak responses (15 frames, 5 s) of cAMP and dopamine during the presentations of shock, OCT, MCH, and ACV. (B, C) Two-photon live imaging on the γ1-γ5 compartments of KCs in genetic control flies (B) and *Dop1R1^KO^* mutant flies (C) expressing jRGECO1b and G-Flamp1 under the control of the γCRE-p-GAL4 driver. Averaged calcium responses during a 40-second air exposure were used to show ROIs for each compartment, depicted in grayscale. ΔF/F0 images of averaged peak responses (15 frames, 5 s) of calcium and cAMP during the presentations of shock, OCT, MCH, and ACV. (D–F) The averaged peak ΔF/F0 responses of cAMP and dopamine (D), and Ca^2+^ and cAMP (E and F), were quantified for each compartment. (D) Quantification of the responses across flies, including the representative sample shown in (A), using the same 15-frame (5-s) averaging window for each sample. Data from different compartments obtained from the same fly are connected by dashed lines. Bar graphs represent mean values ± SEM. For (D), γ1, n = 6 and γ2–γ5, n = 10; for (E), γ1, n = 9 and γ2–γ5, n = 10; and for (F), γ1, n = 6 and γ2–γ5, n = 8. Within each stimulus condition (OCT, MCH, ACV), responses in the γ2–γ5 compartments were compared using the Friedman test followed by Dunn’s multiple comparisons test for all pairwise comparisons. Exact p-values are shown where significant differences were observed. Values for the γ1 compartment are shown for reference only and were excluded from the statistical analyses because γ1 measurements were not obtained from all flies. (G) The changes in cAMP (red) and dopamine (green) levels over 15–30 s in response to different odors (OCT, MCH, and ACV) in five γ lobe compartments (γ1-γ5). The odors were presented by opening the solenoid valve between 15-18 seconds. Solid lines represent the average response from 7 flies, and shaded areas show the standard error of the mean (SEM). The datasets used for the analyses in (D) and (G) were obtained from independent cohorts of flies; therefore, the sample sizes differ between the two panels. (H) Comparison of the odor-evoked calcium responses in the γ2-γ5 compartments between control and *Dop1R1^KO^* mutant flies. Data are presented as mean ± SEM. p-values are shown for comparisons where a significant difference was observed. Mann-Whitney U test, control: n = 10; *Dop1R1^KO^*: n = 8. Figure 2A-video and Figure 2D-video show simultaneous cAMP and dopamine imaging, whereas Figure 2B-video, Figure 2C-video, Figure 2E-video, and Figure 2F-video show simultaneous calcium and cAMP imaging. All videos were acquired at 3 Hz and played back at 9 Hz. Figure 2A-video (related to Figure 2A and D). Figure 2B-video (related to Figure 2B, E and H). Figure 2C-video (related to Figure 2C and H). Figure 2D-video (related to Figure 2D). Figure 2E-video (related to Figure 2E and H). Figure 2F-video (related to Figure 2F and H).

### cAMP levels are not tightly coupled to calcium during odor–shock pairing

As previously noted, the long-held principle of coincidence detection posits that cAMP levels in CS+ responsive KCs should be specifically elevated when the odor stimulus and electric shock are paired. How does this occur, despite the fact that the odor-evoked cAMP rise is not tightly coupled to local calcium concentrations but instead depends predominantly on dopamine, which in the γ1-2 compartments is released at least in part by PPL1 DANs that also convey shock information? To answer this question, we conducted imaging of calcium and cAMP transients in the γ lobe of head-fixed flies that expressed G-Flamp1 and jRGECO1b driven by the γCRE-p driver and were subjected to OCT in conjunction with the delivery of electric shock, mimicking the established paradigm of aversive memory conditioning. Electric shock increased cAMP levels in the γ1-2 compartments (Figure 1E), as two sets of PPL1 DANs, γ1pedc and γ2α′1, are known to convey aversive signals. We examined the cAMP transients specifically in the γ1 and γ2 compartments in an enlarged imaging field. A substantial increase in cAMP was observed across the entirety of each compartment irrespective of localized calcium elevations, two examples of which are shown in Figure 3A. In both the γ1 and γ2 compartments, the mean response to paired OCT–shock stimulation did not exceed the sum of the mean responses to each stimulus presented alone (Figure 3B). In the γ2 compartment, the mean paired response was numerically lower than this sum. These results were not unexpected, as the cAMP levels depend largely on the dopamine release from the same DANs, which are activated by both CS and US. However, these findings differ from a previous report of a synergistic (supra-additive) increase in cAMP upon pairing (Tomchik and Davis, 2009), which was obtained by bath application of acetylcholine and dopamine to ex vivo brain explants and in the α/β and α′/β′ lobes. Notably, as reported previously (Honegger et al., 2011), approximately 5–9% of KCs exhibit odor-evoked calcium responses, whereas in our data, the majority of KCs exhibit cAMP synthesis in response to dopaminergic input during olfactory stimulation. Thus, some overlap between calcium and cAMP signals is plausible, even without a direct causal relationship. Indeed, such overlap was observed. However, it is crucial to determine the extent to which the two signals are correlated across the entire field of the compartment.

**Figure 3.**
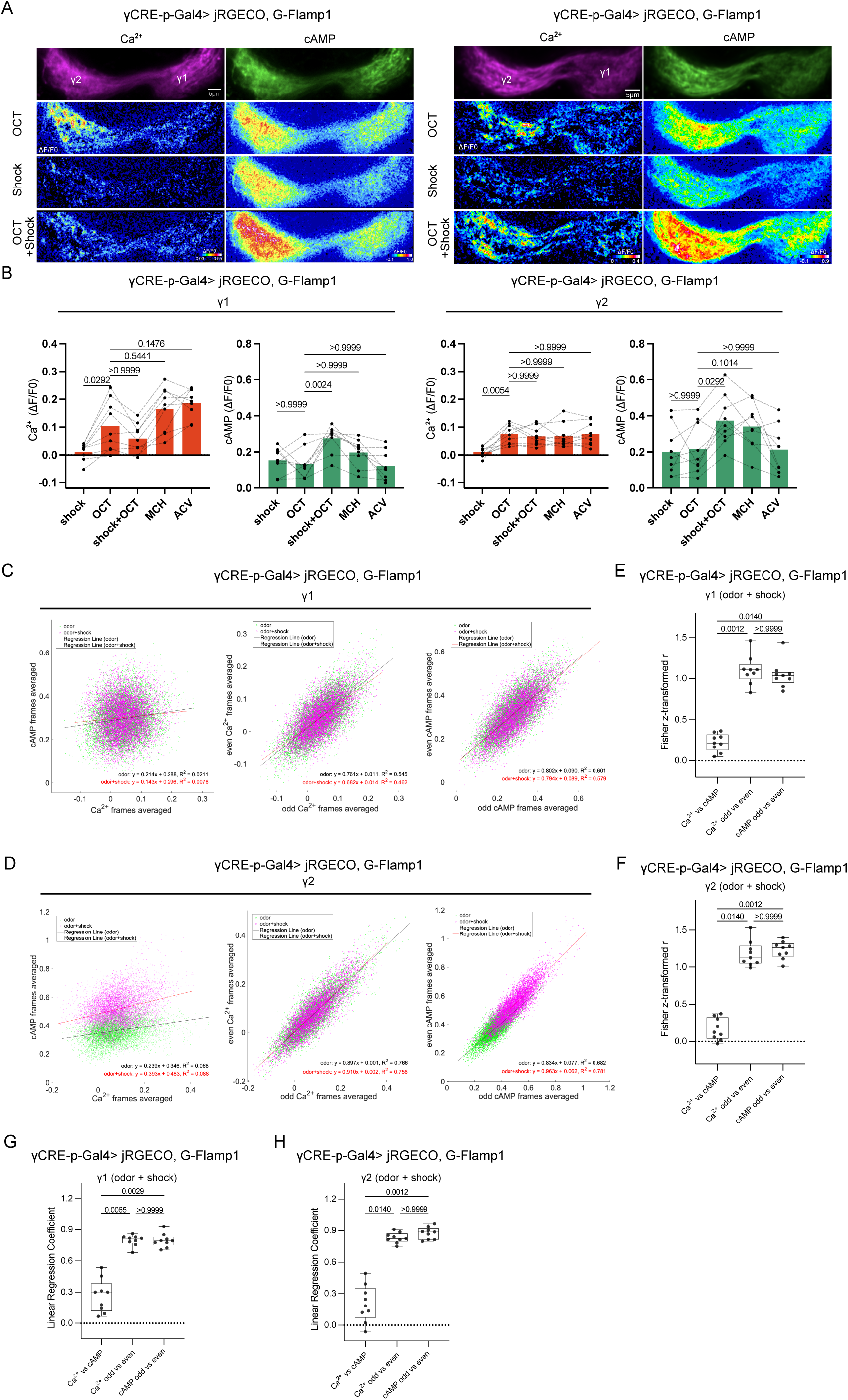
cAMP levels are not tightly coupled to calcium during odor–shock pairing. (A) The top panels display the averaged representations of raw intensity values extracted from 80 frames, spanning a duration of 40 seconds including both stimulus and non-stimulus periods. The three lower rows show representative ΔF/F0 images of averaged peak calcium and cAMP responses (10 frames over 5 s) to OCT alone, shock alone, and paired OCT– shock stimulation in the γ1 and γ2 compartments of flies expressing jRGECO1b and G-Flamp1 under the control of the γCRE-p driver. (B) The averaged peak ΔF/F0 responses were quantified. Data obtained under different stimulus conditions from the same fly are connected by dashed lines. Bar graphs represent mean values ± SEM. The stimulus conditions were compared using the Friedman test followed by Dunn’s multiple comparisons test, with OCT used as the reference condition. Exact p-values are shown for comparisons of OCT with shock, OCT plus shock, MCH, and ACV. n = 9. (C) Pixel-wise scatter plots of mean ΔF/F0 values from representative γ1 compartment images. Left, calcium versus cAMP for a single animal. Middle, calcium self-correlation control, comparing odd-versus even-frame mean ΔF/F0 values from the same recording. Right, cAMP self-correlation control, comparing odd-versus even-frame mean ΔF/F0 values. All panels use the same dataset as in (B). Linear fits are overlaid. OCT (CS), green; OCT + shock (CS + US), magenta. In this representative γ1 dataset, the raw correlation coefficient for Ca^2+^ versus cAMP during paired OCT–shock stimulation was r = 0.0872, whereas the corresponding self-correlation coefficients were r = 0.679 for Ca^2+^ and r = 0.760 for cAMP. (D) Same as (C), for the γ2 compartment. Left, calcium versus cAMP; middle, calcium self-correlation control; right, cAMP self-correlation control. In this representative γ2 dataset, the raw correlation coefficient for Ca^2+^ versus cAMP was r = 0.297, whereas the corresponding self-correlation coefficients were r = 0.869 for Ca^2+^ and r = 0.883 for cAMP. (E) Quantification of Fisher’s z-transformed correlation coefficients in γ1. Three measures were compared: Ca^2+^ versus cAMP, Ca^2+^ odd-versus-even self-correlation, and cAMP odd-versus-even self-correlation. A significant overall difference was detected by Friedman test (p = 0.0002). Post hoc multiple-comparison analysis showed that Ca^2+^–cAMP correlations were significantly lower than Ca^2+^ odd-versus-even self-correlations (p = 0.0012) and cAMP odd-versus-even self-correlations (p = 0.0140), whereas no significant difference was detected between the two self-correlation measures (p > 0.9999). (F) Quantification of Fisher’s z-transformed correlation coefficients in γ2. Three measures were compared as in (E). A significant overall difference was detected by Friedman test (p = 0.0002). Post hoc multiple-comparison analysis showed that Ca^2+^–cAMP correlations were significantly lower than Ca^2+^ odd-versus-even self-correlations (p = 0.0140) and cAMP odd-versus-even self-correlations (p = 0.0012), whereas no significant difference was detected between the two self-correlation measures (p > 0.9999). (G) Quantification of regression coefficients in γ1 using the same datasets as in (E). A significant overall difference was detected by Friedman test (p = 0.0003). Post hoc multiple-comparison analysis showed that regression coefficients for Ca^2+^ versus cAMP were significantly lower than those for Ca^2+^ odd-versus-even self-correlation (p = 0.0065) and cAMP odd-versus-even self-correlation (p = 0.0029), whereas no significant difference was detected between the two self-correlation conditions (p > 0.9999). (H) Quantification of regression coefficients in γ2 using the same datasets as in (F). A significant overall difference was detected by Friedman test (p = 0.0002). Post hoc multiple-comparison analysis showed that regression coefficients for Ca^2+^ versus cAMP were significantly lower than those for Ca^2+^ odd-versus-even self-correlation (p = 0.0140) and cAMP odd-versus-even self-correlation (p = 0.0012), whereas no significant difference was detected between the two self-correlation conditions (p > 0.9999). Panels E–H are based on the OCT + shock trials. Odor–shock pairing was repeated a variable number of times across samples; only the first paired trial in each series was used for quantification. n = 9 for panels E–H. Figure 3A-video (related to Figure 3A-F). Figure 3B-video (related to Figure 3B-F). Videos illustrating simultaneous calcium and cAMP imaging, acquired at 2 Hz and played back at 6 Hz.

The relationship between calcium and cAMP levels was further visualized using scatter plots: γ1 was analyzed using data from the left panel of Figure 3A (Figure 3C, left), and γ2 using data from the right panel of Figure 3A (Figure 3D, left). In the γ1 and γ2 example datasets, the pixel-wise correlation between Ca^2+^ and cAMP signals during paired OCT–shock stimulation was low, with raw correlation coefficients of r = 0.0872 in γ1 and r = 0.297 in γ2 (Figures 3C and 3D). To determine whether this weak Ca^2+^–cAMP correspondence was consistent across samples, we calculated correlation coefficients for each sample in γ1 and γ2. The mean correlation coefficients, obtained by averaging Fisher’s z-transformed values and then back-transforming them to r values, were also low for Ca^2+^ versus cAMP in both γ1 and γ2 (r = 0.219 and 0.166, respectively). Although these values should not be interpreted as the mean R² across samples, they correspond to squared back-transformed mean r values of approximately 0.048 and 0.028 in γ1 and γ2, respectively, further supporting the view that the pixel-wise correspondence between Ca^2+^ and cAMP signals was limited in both compartments.

We next compared Fisher’s z-transformed correlation coefficients across three measures: Ca^2+^ versus cAMP, Ca^2+^ odd-versus-even self-correlation, and cAMP odd-versus-even self-correlation. The within-signal self-correlations were used as empirical internal benchmarks for the degree of correspondence detectable under the present imaging conditions (Figures 3E and 3F). In both γ1 and γ2, Fisher’s z-transformed Ca^2+^–cAMP correlation coefficients were significantly lower than those for either Ca^2+^ or cAMP self-correlation, whereas no significant difference was detected between the two self-correlation measures.

Together, these analyses indicate that, in both γ1 and γ2, Ca^2+^ and cAMP signals showed only weak spatial correspondence, and that this weak correspondence was substantially lower than the within-signal correspondence observed under the same imaging conditions. Because the self-correlation values of the Ca^2+^ and cAMP signals were comparably high in both compartments, the low Ca^2+^–cAMP correlation is difficult to explain simply by reduced reproducibility in only one of the two signals. Rather, the data suggest that although calcium may contribute in part to cAMP production, the spatial pattern of cAMP cannot be accounted for solely by the spatial distribution of calcium.

Accordingly, this weak spatial correspondence is not readily explained by a simple model in which pairing with electric shock selectively elevates cAMP in local regions of the γ compartment that exhibit large odor-evoked Ca^2+^ responses. The observed cAMP levels were not entirely contingent upon calcium, even under paired odor and electric shock stimulation. This conclusion should nevertheless be interpreted cautiously, because the present analysis does not exclude the possibility that calcium contributes partially or indirectly to cAMP synthesis, nor does it rule out localized coincidence-detection processes that may not have been fully resolved under the present imaging conditions.

As a complementary analysis, we also compared the slopes of the regression lines derived from the same datasets in both compartments (Figures 3G and 3H). This analysis yielded a pattern similar to that of the correlation analysis: in both γ1 and γ2, the regression coefficients for Ca^2+^ versus cAMP were lower than those for the self-correlation controls, whereas no significant difference was detected between the two self-correlation conditions. Because regression coefficients are influenced by signal scaling, we treated the comparison of Fisher’s z-transformed correlation coefficients as the primary statistical analysis. Taken together, these two analyses support the interpretation that the correspondence between Ca^2+^ and cAMP signals was weak and was lower than that observed between repeated measurements within each signal in both compartments. These findings therefore suggest that cAMP concentration alone is unlikely to be sufficient to distinguish the KCs that respond to the CS+ odor from the rest of the KC population.

### Aversive conditioning shifts the cAMP code evoked by the CS+ toward a negative-valence state

As suggested above, the cAMP level in the γ1-2 compartments may depend on dopamine released by PPL1 DANs (Aso et al., 2012), whereas those in the γ3-γ5 compartments may depend on dopamine release by PAM DANs (Liu et al., 2012; Yamagata et al., 2016). Moreover, discrete compartments can be characterized in relation to odor valence: activity in DANs innervating the γ2 and γ4 compartments is associated with aversive and appetitive olfactory valence, respectively (Kato et al., 2023; Siju et al., 2020). Furthermore, DANs integrate odor valence and shock signals (Huang et al., 2024). Given that cAMP may act as an intermediary of dopaminergic signaling in determining odor valence, its spatial distribution across γ-lobe compartments of KCs may constitute a representation of integrated valence. We refer to this compartmental pattern of cAMP levels across the γ lobe as the “cAMP code” (Figure 10B). We therefore hypothesized that aversive conditioning would reshape the cAMP code toward a negative-valence-associated state. Although the absolute cAMP levels varied among individuals for each odor, we reasoned that the relative cAMP levels of the γ2 (corresponding to negative valence) and γ4 (corresponding to positive valence) compartments, the γ2/γ4 cAMP ratio, could serve as a metric for evaluating the valence of an olfactory stimulus for an individual. As such, changes in the γ2/γ4 cAMP ratio prior to and post-conditioning could be employed to assess the differential response to the stimulus, and may therefore reflect the acquisition of memory. To this end, we attempted to recapitulate aversive conditioning by subjecting a fly to three consecutive episodes of electric shocks while exposing it to OCT, followed by exposure to MCH and ACV without electric shocks (OCT as CS+), or exposing it to MCH with electric shocks and to OCT and ACV without electric shocks (MCH as CS+), while observing it under a microscope (Figure 4A). During the process, cAMP transients were recorded for each exposure and quantified. Averaged images of a single sample and profiles of cAMP transients from pooled samples are presented (Figure 4B and E). Note that, based on the innate behavioral responses, OCT and MCH are both similarly aversive compared to air, whereas ACV is attractive (Badel et al., 2016). Electric shock elicited elevated cAMP levels in the γ1–γ3 compartments. Because the γ1 compartment often lay in a different focal plane from the other compartments, γ1 data could not be obtained from all samples; γ1 was therefore excluded from the statistical analyses in this and subsequent quantifications. During the conditioning protocol in which OCT was paired with electric shock, cAMP levels in the γ2 compartment tended to increase, consistent with the results in Figure 3, although this change did not reach statistical significance in the present experimental series (Figure 4B, E and K). This was in parallel with a significant decrease in cAMP levels in the γ4 compartment. After conditioning, the presentation of OCT reverted cAMP levels to pre-conditioning states in the γ2 compartment, while maintaining lower cAMP levels in the γ4 and γ5 compartments relative to pre-conditioning states (Figure 4H). Accordingly, the γ2/γ4 cAMP ratio for OCT increased from Pre to Post, whereas no significant changes were detected for MCH or ACV, neither of which had been paired with an electric shock (Figure 4N). Similarly, when MCH was paired with an electric shock (Figure 4C, F and I), there was an increase in the γ2/γ4 cAMP ratio in response to MCH (p = 0.0078), albeit to a lesser degree, while no significant changes were observed in either OCT or ACV (Figure 4O), even though γ4 responses to these CS− odors also decreased by Cond.3 relative to Pre (OCT, p = 0.0150; ACV, p = 0.0030; Figure 4L). Thus, the persistent shift followed the odor paired with shock rather than odor identity.

**Figure 4.**
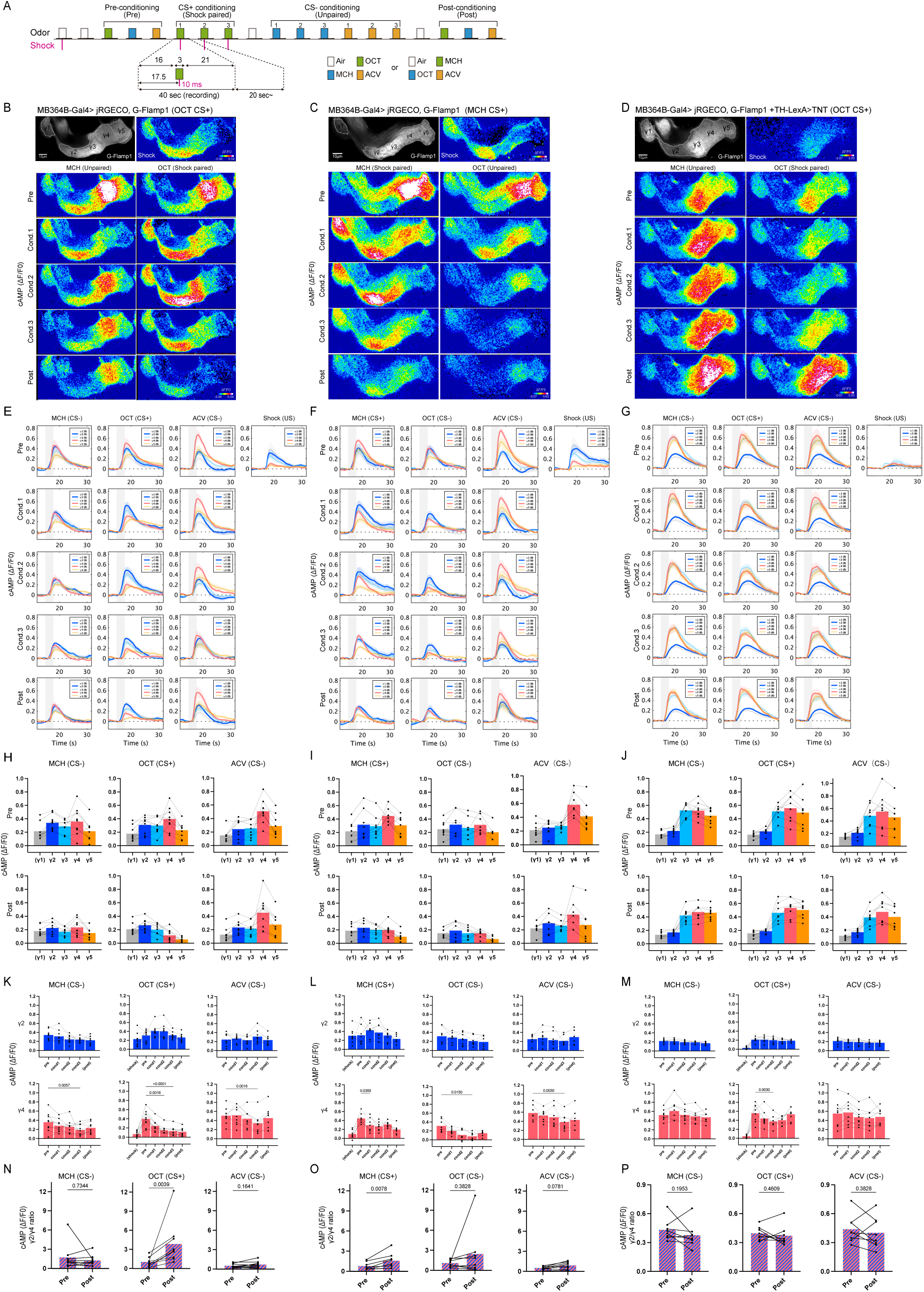
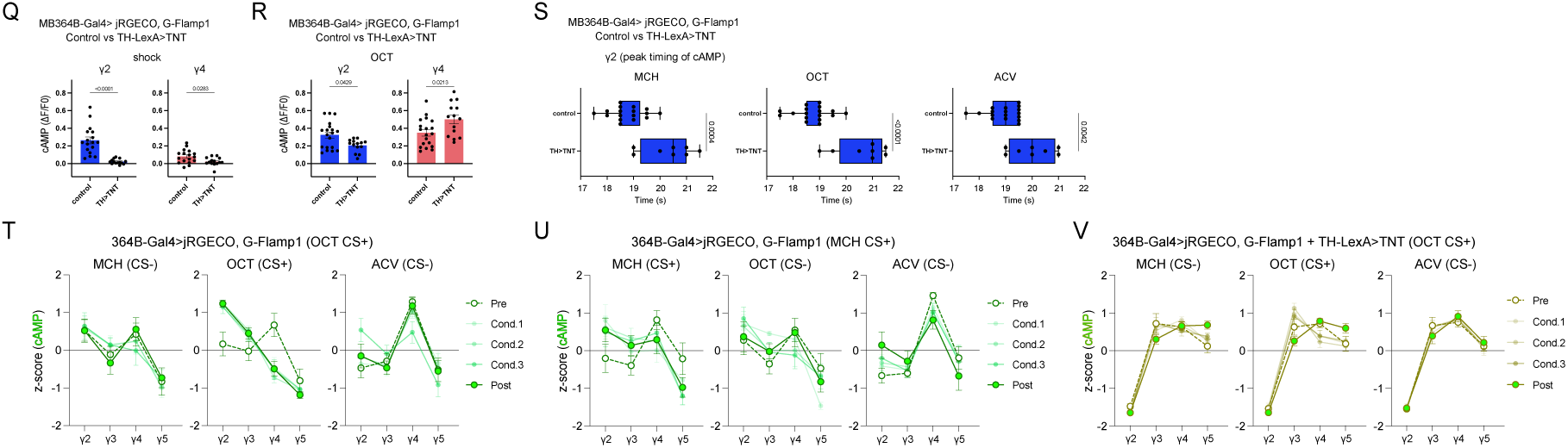
Aversive conditioning shifts the cAMP code evoked by the CS+ toward a negative-valence state. (A) Schematic representation of conditioning procedure. Sixteen seconds after the start of the recording, a 3 second odor stimulus was presented to the fly. Following a delay of 1.5 seconds from the onset of the odor presentation, an electric shock of 60 volts was delivered to the fly for a duration of 10 milliseconds. In one series of experiments, OCT was paired with electric shock, while MCH and ACV were not (B and D). In another set of experiments, MCH was paired with electric shock, while OCT and ACV were not (C). (B, C, D) All the cAMP images in the γ1-γ5 compartments of KCs were captured from flies expressing G-Flamp1 and jRGECO1b under the control of the MB364B driver. An averaged raw intensity image of cAMP derived from 80 frames acquired during air presentation for 40 s, onto which the ROIs for each compartment are mapped (upper left) and the ΔF/F0 image obtained by averaging the responses (5 s, 10 frames) to electric shock (upper right) are shown. The images displayed below depict the ΔF/F0 images obtained by averaging the responses (5 s, 10 frames) to OCT (as CS+) during the pre-conditioning, first, second, and third pairings, as well as post-conditioning. Additionally, the images illustrate the neural responses to MCH (as CS−) during the same experimental procedure, but without electric shock (B). Similarly, the corresponding images obtained when MCH was presented as the CS+ and OCT as the CS− are shown in (C). In (D), TNT was expressed in PPL1 DANs by the TH-LexA driver to block neurotransmitter release from their terminals in the compartments they innervate. OCT was paired with electric shock, while MCH and ACV were not, following the same procedure as in (B). (E, F, G) A series of line plots depicting the temporal dynamics of cAMP over time in seconds (s) across different stimulus conditions. Each subplot corresponds to a specific stimulus and experimental stage. Each plot shows the ΔF/F0 of cAMP of the γ1-γ5 compartments. The solid lines represent the mean values for each compartment, calculated from the number of flies indicated in parentheses within the subplots. The shaded areas around each solid line represent the SEM. Shaded rectangles indicate the estimated period of odor delivery to the fly (approximately 16–19 s). A 60-V electric shock was delivered at 17.5 s for 10 ms. (H, I, J) The mean peak ΔF/F0 (10 frames for 5 s) values of each compartment were averaged and represented in bar graphs, generating cAMP codes for the pre- and post-conditioning responses to each odor. The plots of different compartments derived from the same sample are connected with dashed lines. n = 9 (H), 8 (I), and 8 (J). (K–M) Changes in cAMP responses in the γ2 and γ4 compartments across the conditioning steps were analyzed. For each compartment, the analyzed responses were compared using the Friedman test followed by Dunn’s multiple comparisons test, with the corresponding pre-conditioning odor response used as the reference condition and compared with Cond.1–3. The same datasets used in (H), (I), and (J) were used for (K), (L), and (M), respectively. Shock-evoked responses during CS+ conditioning and post-conditioning responses to each stimulus are shown for reference but were not included in the statistical analyses. Exact p-values are shown where significant differences were observed. n = 9 for (K), n = 8 for (L), and n = 8 for (M). (N, O, P) The ratio of cAMP levels in the γ2 compartment to those in the γ4 compartment for each odor response was calculated. Bar graphs represent mean values. The ratios obtained before and after the conditioning phase are then compared by Wilcoxon signed rank test. The same datasets used in (H), (I), and (J) were used for (N), (O), and (P), respectively. p-values are shown for all comparisons. n = 9 (N), 8 (O), and 8 (P). (Q, R) Shock-evoked (Q) and OCT-evoked (R) cAMP responses in the γ2 and γ4 compartments of control and TH-LexA>TNT flies. Bars represent mean ± SEM. Data include pre-conditioning responses of flies subjected to the conditioning paradigm and responses of additional flies recorded without conditioning. Mann–Whitney U test; (Q) control, n = 17; TH-LexA>TNT, n = 14; (R) control, n = 19; TH-LexA>TNT, n = 14. (S) Peak timing of odor-evoked cAMP responses in the γ2 compartment, determined at the frame resolution (0.5 s). For TH-LexA>TNT, only flies subjected to the conditioning paradigm were analyzed in (S). Box plots show the median, interquartile range, and minimum–maximum range, with individual data points overlaid. Mann–Whitney U test; control, n = 17; TH-LexA>TNT, n = 8. (T-V) cAMP profile (γ2-γ5) across Pre, Cond.1–3, and Post for OCT, MCH, and ACV application side by side for three fly populations: flies in which OCT was paired with shock (n = 9; T), flies in which MCH was paired with shock (n = 8; U), and flies expressing TNT in PPL1 dopaminergic neurons (TH-LexA>TNT; OCT paired with shock; n = 8; V). Each panel shows mean ± SEM at each phase; corresponding phase labels are shown alongside each panel. Figure 4B-video (related to Figure 4B, E, H, K and N). Figure 4C-video (related to Figure 4C, F, I, L and O). Figure 4D-video (related to Figure 4D, G, J, M and P-S). Figure 4E-video (related to Figure 4E, H, K and N). Figure 4F-video (related to Figure 4F, I, L and O). Figure 4G-video (related to Figure 4G, J, M and P-S). Videos illustrating cAMP imaging, acquired at 2 Hz and played back at 6 Hz.

We next conducted the conditioning experiment with flies in which PPL1 DANs were silenced through the expression of the tetanus toxin light chain (Sweeney et al., 1995), utilizing the TH-LexA > LexAop-TNT system (Karuppudurai et al., 2014)(Figure 4D, G, J, M and P). DANs located within the PPL1 cluster are known for transmitting aversive information to the γ1 and γ2 compartments. Consistent with this, shock-evoked cAMP responses in the γ2 compartment were nearly absent in these flies (Figure 4Q). Odor-evoked cAMP responses, however, were still detected, possibly reflecting other signals (Figure 4R). Note that the peak of these cAMP responses was delayed by approximately 1– 2 s compared to the control (Figure 4S). Although γ4 responses to OCT decreased during pairing even in these flies (Figure 4M), they returned to near pre-conditioning levels after conditioning (Figure 4J), and the γ2/γ4 cAMP ratio did not change from Pre to Post for any odor (Figure 4P). Thus, the γ4 decrease during pairing can occur even when PPL1 output is blocked, whereas PPL1 output may be needed for its persistence and the resulting shift in the γ2/γ4 balance. Interestingly, odor-evoked cAMP responses in the γ4 compartment were elevated in flies with silenced PPL1 DANs compared with controls (Figure 4R), and those in γ3 and γ5 showed a similar tendency (Figure 4H, J). Considering the projection of PAM-class dopamine neurons into the γ3-γ5 compartments, this finding suggests a reciprocal regulatory mechanism between PPL1 DANs and PAM DANs. It further points to a circuit-level mechanism that may coordinate activity across these two dopaminergic populations. Such coordination may involve interneurons such as CRE011, which integrates input from nine types of MBONs, including γ2α′1, and provides input to reward DANs, including those innervating the γ4 compartment (Shuai et al., 2024).

To visualize how the compartmental cAMP code evolves during conditioning, we plotted, for each fly population, the within-sample z-scored cAMP profile (γ2-γ5) across the phases of conditioning (Pre, Cond.1-3, Post; Figure 4T-V). For flies in which OCT served as the reinforced odor (CS+; Figure 4T), the OCT profile shifted markedly between Pre and Post, with relative cAMP levels increasing in γ2 and decreasing in γ4 (γ2-γ4 difference: Pre = −0.50, Post = +1.73). A qualitatively similar shift was observed in flies for which MCH instead served as the reinforced odor (Figure 4U; γ2–γ4 difference: Pre = −1.02, Post = +0.25). In flies expressing TNT in PPL1 dopaminergic neurons (TH-LexA>TNT), the cAMP profile was essentially indistinguishable across the three odors tested (Figure 4V), in contrast to the clearly odor-dependent Pre profiles observed when PPL1 neurotransmission was intact (Figure 4T and 4U). This loss of odor-specific patterning was accompanied by the absence of a persistent conditioning-associated shift. Together, these results again suggest that intact PPL1 neurotransmission contributes to both the odor-specific compartmentalization of the cAMP code and its conditioning-associated reorganization.

### Aversive conditioning induces compartmental changes in dopamine levels that are reflected in the cAMP code

Given that the cAMP code is shaped by dopaminergic signaling, we predicted that cAMP dynamics would closely parallel those of dopamine during aversive conditioning. Flies expressing both the GRAB_DA2m and Pink Flamindo sensors, driven by the γCRE-p GAL4 driver, were subjected to aversive conditioning while being imaged under a microscope. Averaged images and time courses of ΔF/F0 for cAMP and dopamine levels are presented (Figures 5A–B′). The two signals showed broadly similar dynamics throughout the conditioning process, although dopamine levels rose earlier and more abruptly than cAMP levels (Figures 5A–C′). Dopamine levels in the γ2 compartment during conditioning tended to be higher than the pre-conditioning odor responses, although the difference was not significant (Figure 5C and D). Conversely, dopamine levels in the γ4 compartment decreased significantly relative to the Pre condition (Figure 5C and D). These alterations were reflected in the concurrently recorded cAMP levels (Figure 5C′ and D′). This suggests that the changes in cAMP levels during conditioning largely reflect the changes in dopamine levels, without requiring a synergistic interaction between dopamine and calcium at the Rutabaga adenylyl cyclase. cAMP responses to ACV (CS−) in γ4 also decreased during conditioning (Figure 5D′), although no corresponding significant decrease in dopamine was detected (Figure 5D). Following conditioning, the dopamine γ2/γ4 ratio in response to OCT, the CS+ odor, was markedly higher than that observed before conditioning (Figure 5E). This increase was paralleled by a corresponding increase in the cAMP γ2/γ4 ratio (Figure 5E′).

**Figure 5.**
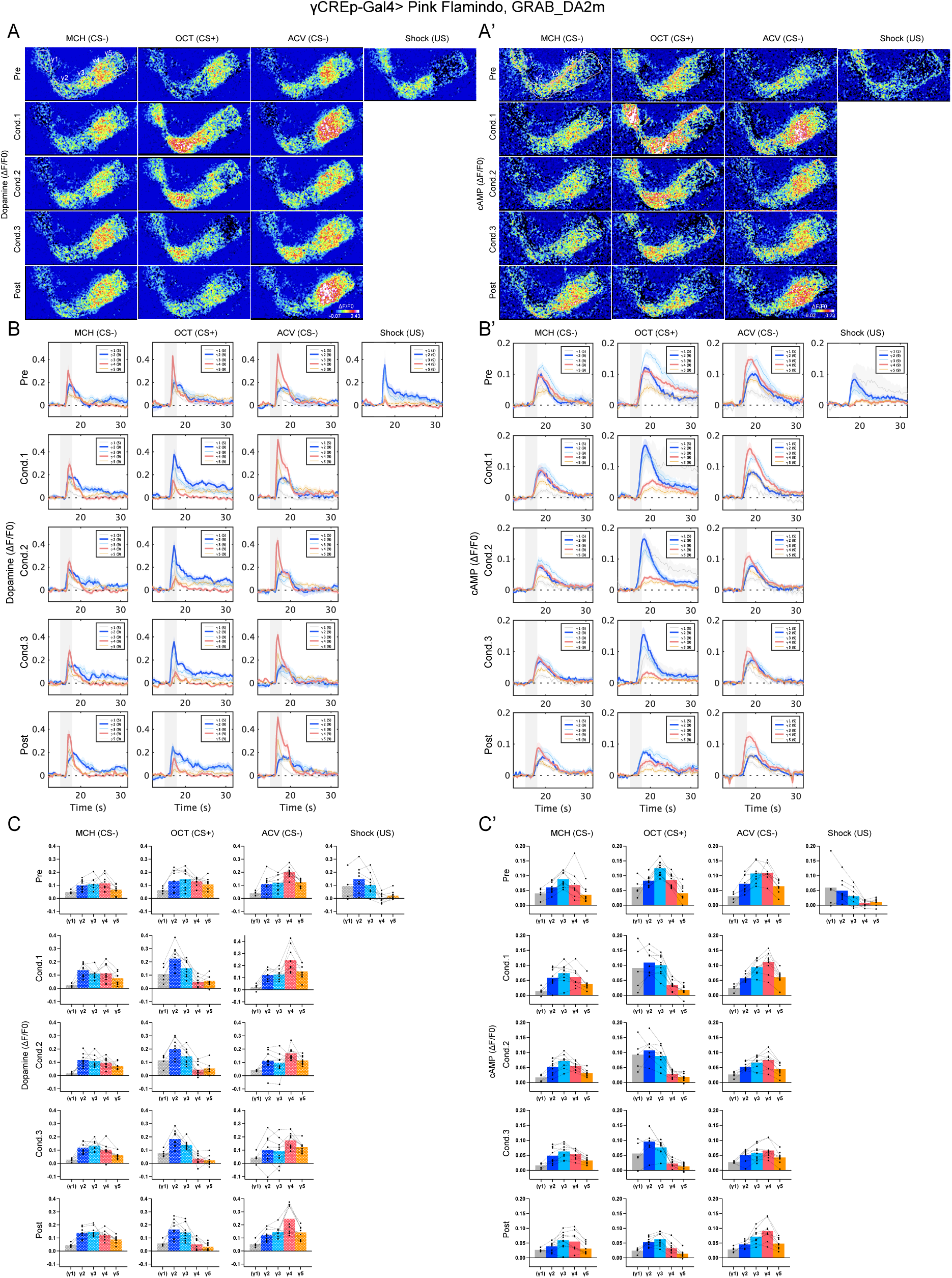
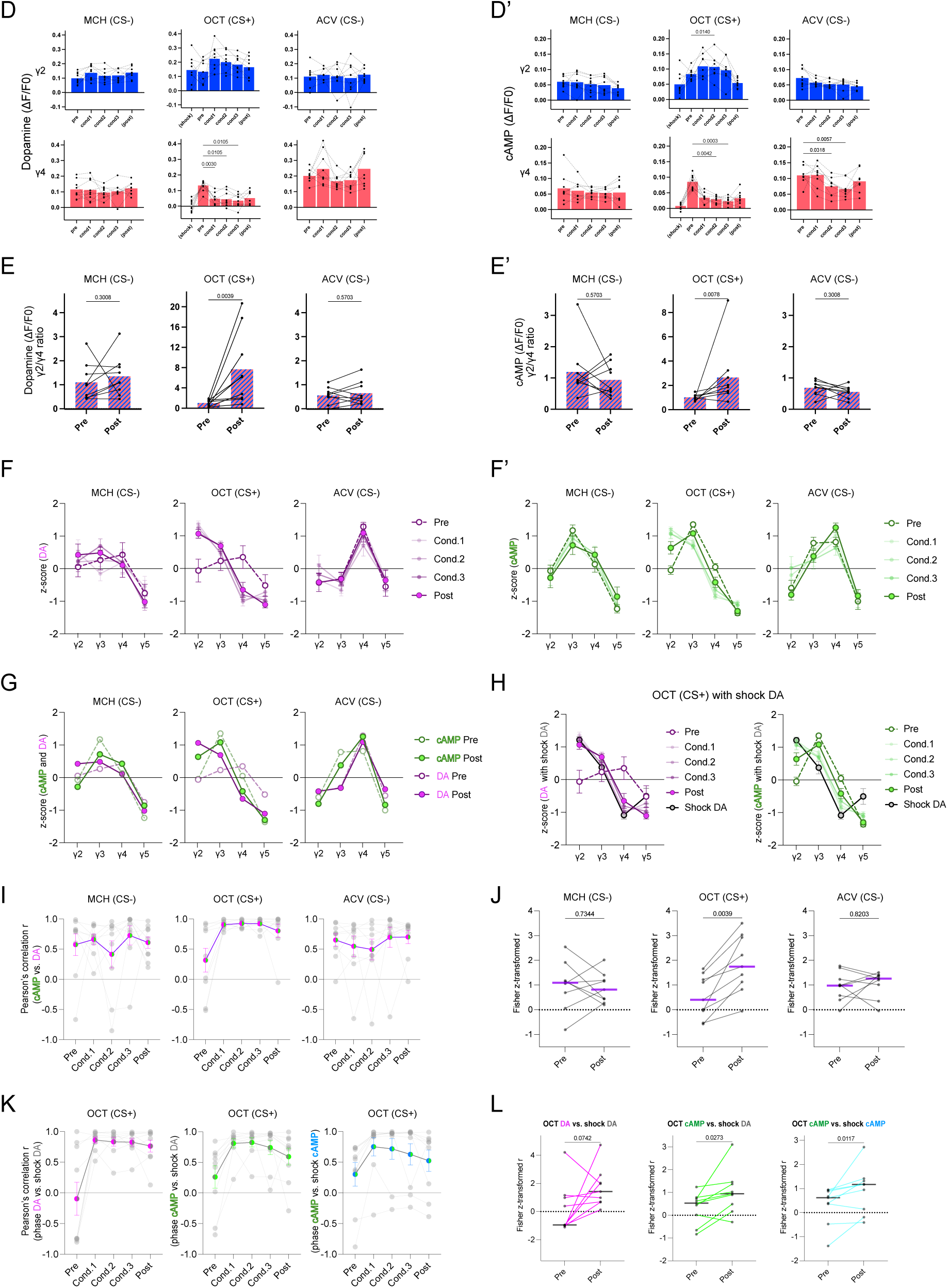
Aversive conditioning induces compartmental changes in dopamine levels that are reflected in the cAMP code. (A, A′) The ΔF/F0 images, depicting the averaged peak responses (15 frames for 5 s) of Dopamine (A) and cAMP (A′) in the γ1-γ5 compartments of KCs, were captured from flies expressing Pink Flamindo and GRAB_DA2m sensors under the control of the γCRE-p driver during the presentations of OCT (CS+), MCH (CS−), ACV (CS−), and shock (US) throughout conditioning. The ROIs for each compartment are mapped onto the top left panel. (B, B′) A series of line plots depicting the temporal dynamics of Dopamine (B) and cAMP (B′) over time in seconds (s) across different stimulus conditions. Each subplot corresponds to a specific stimulus and experimental stage. The y-axis indicates ΔF/F0. The solid lines represent the mean values for each compartment, calculated from the number of flies indicated in parentheses within the subplots. The shaded areas around each line represent the SEM. Shaded rectangles indicate the estimated period of odor delivery to the fly (approximately 16–19 s). A 60-V electric shock was delivered at 17.5 s for 10 ms. (C, C′) Quantification of the averaged peak ΔF/F0 responses during the pre-conditioning, Cond. 1–3, and post-conditioning stages, together with the shock-evoked responses. The plots of different compartments derived from the same sample are connected with dashed lines. The bar graph illustrates the mean values. (D, D′) Dopamine (D) and cAMP (D′) responses in the γ2 and γ4 compartments were compared using the Friedman test followed by Dunn’s multiple comparisons test, with the corresponding pre-conditioning odor response used as the reference condition. The same datasets used in (C) and (C′) were used for (D) and (D′), respectively. Shock-evoked responses in the CS+ experimental group, as well as post-conditioning responses to each stimulus, are shown for reference but were not included in the statistical analyses. p-values are shown for comparisons with significant differences. n = 9 for all stimulus sets in both γ2 and γ4. (E, E′) Comparisons of γ2/γ4 dopamine ratios before and after conditioning for each odor (E), and γ2/γ4 cAMP ratios before and after conditioning (E′). The same datasets used in (C) and (C′) were used for (E) and (E′), respectively. p-values are shown for all comparisons. n = 9, 9, 9 (E), and 9, 9, 9 (E′) for MCH, OCT, and ACV, respectively. Wilcoxon signed-rank test. (F, F′) Within-sample z-scored DA (F) and cAMP (F′) profiles (γ2–γ5) evoked by MCH (CS−), OCT (CS+), and ACV (CS−) across Pre, Cond.1–3, and Post. Mean ± SEM at each phase; n = 9. (G) For each odor, the Pre and Post profiles of DA (F) and cAMP (F′) are overlaid on the same axes. (H) The group-averaged shock-evoked DA profile (mean ± SEM across flies), obtained by presenting electric shock alone, is overlaid as a fixed reference on the OCT (CS+) DA (left) and cAMP (right) profiles shown in F and F′. (I) Pattern correlation r between DA and cAMP profiles across phases for MCH, OCT, and ACV. Small grey circles show individual flies; colored circles and error bars show mean ± SEM. (J) Fisher z-transformed pattern correlation between DA and cAMP at Pre and Post for each odor. Lines connect data from the same fly; horizontal bars indicate the median. p-values are shown for all comparisons (two-sided Wilcoxon signed-rank test). (K) Pattern correlation r between the OCT-evoked profile and each fly’s own shock-evoked profile across phases: DA vs. shock-evoked DA (left), cAMP vs. shock-evoked DA (middle), and cAMP vs. shock-evoked cAMP (right). Small grey circles show individual flies; colored circles and error bars show mean ± SEM. (L) Fisher z-transformed pattern correlations in K at Pre and Post. Lines connect data from the same fly; horizontal bars indicate the median. p-values are shown for all comparisons (two-sided Wilcoxon signed-rank test). Figure 5AA′-video (related to Figure 5AA′-EE′). Figure 5BB′-video (related to Figure 5BB′-EE′). Videos illustrating simultaneous dopamine and cAMP imaging, acquired at 3 Hz and played back at 9 Hz.

To directly visualize this compartmental reorganization, we plotted each signal’s within-sample z-scored profile (γ2–γ5) across the phases of conditioning for each of the three odors tested. For OCT (CS+), DA showed a pronounced shift that emerged at Cond.1 and was sustained through Post: relative activity increased in γ2 and decreased in γ4, whereas γ3 and γ5 changed comparatively little (Figure 5F). cAMP showed a qualitatively similar shift in the same two compartments, although its Pre profile already carried a prominent γ3 peak that was retained throughout conditioning (Figure 5F′). To ask how closely the cAMP profile tracks that of DA, we overlaid the Pre and Post DA and cAMP profiles for each odor (Figure 5G). We then quantified the similarity using a pattern correlation (r): the Pearson correlation between the two signals’ relative levels across the four γ-compartments (γ2–γ5) within the same fly, after normalizing out each fly’s overall signal amplitude (Figure 5I). A high positive r indicates that the two signals have similar relative values across compartments, an r near zero indicates little linear relationship between their compartmental distributions, and a negative r indicates a mirror-image relationship. For OCT (CS+), mean r rose from 0.32 ± 0.20 at Pre to 0.91 ± 0.03, 0.93 ± 0.02, and 0.92 ± 0.03 during Cond.1, Cond.2, and Cond.3, respectively, and remained elevated at 0.80 ± 0.11 at Post (Figure 5I). Because responses during Cond.1–3 include a shock-evoked component, we restricted statistical testing to the comparison between Pre and Post, both of which were evoked by odor alone. The Fisher z-transformed DA–cAMP pattern correlation increased from Pre to Post for OCT (p = 0.0039), but not for MCH (p = 0.7344) or ACV (p = 0.8203; Figure 5J). Note that the Pre correlation for OCT was lower than those for MCH and ACV (Figure 5I); after conditioning, the OCT-evoked DA and cAMP profiles reached a degree of alignment comparable to that observed for the CS− odors.

We next asked whether this reorganization brings the odor-evoked profiles closer to the profile evoked by the US itself. We overlaid the group-averaged shock-evoked DA profile (mean ± SEM across flies) onto the within-sample z-scored DA and cAMP profiles for OCT (Figure 5H). For both signals, the Cond.1–Post profiles visually approached the shock-evoked DA template (γ2 = 1.22 ± 0.09, γ3 = 0.37 ± 0.19, γ4 = −1.09 ± 0.12, γ5 = −0.51 ± 0.24), most evidently in γ2 and γ4: whereas the Pre profiles of both DA (γ2 = −0.06, γ4 = 0.35) and cAMP (γ2 = −0.04, γ4 = 0.05) diverged from the shock template at these two compartments, the Cond.1–Post profiles of both signals approached it (DA: γ2 = 1.07–1.35, γ4 = −0.65 to −1.01; cAMP: γ2 = 0.64–1.20, γ4 = −0.42 to −0.86). To quantify this convergence, we correlated, for each fly, the OCT-evoked profile at each phase with that fly’s own shock-evoked profile for three combinations: DA versus shock-evoked DA, cAMP versus shock-evoked DA, and cAMP versus shock-evoked cAMP (Figure 5K, left to right). As expected, the correlations were high during Cond.1–3, when the responses included the shock-evoked component. At Post, when OCT was presented alone, the correlation of the OCT-evoked cAMP profile with both the shock-evoked DA profile (p = 0.0273) and the shock-evoked cAMP profile (p = 0.0117) remained higher than at Pre, and the OCT-evoked DA profile showed a similar trend (p = 0.0742; Figure 5L). Together, these results suggest that conditioning shifts the cAMP code evoked by the CS+ toward the shock-evoked configuration, and that this shift persists in the absence of shock.

### PKA activity broadly reflects the compartmental cAMP pattern

Protein kinase A (PKA) is a major downstream effector of cAMP signaling, making it a useful readout for assessing the functional consequences of the observed cAMP signals. We employed the expression of a PKA sensor ExRai-AKAR2 (Zhang et al., 2021) in combination with Pink Flamindo to examine whether the compartmental pattern of PKA activity coincides with that of cAMP signaling. The changes in PKA activity were more prolonged than those in cAMP signals (Figure 6B). Although the signal-to-noise ratios of these sensors limited our ability to comprehensively analyze signals at the single-cell level, the relative relationship between signal levels in the γ2 and γ4 compartments was broadly preserved between PKA activity and cAMP signals across all odor stimuli tested, as well as in response to electric shock (Figure 6A–C). For each stimulus, we overlaid the within-sample z-scored cAMP and PKA profiles on the same axes (Figure 6D). Across all four stimuli, the two signals showed an overall similar compartmental shape, without any single stimulus standing out as visibly more or less concordant between cAMP and PKA. This impression was supported quantitatively: the mean pattern correlation r between cAMP and PKA exceeded 0.6 for every stimulus (OCT: 0.65 ± 0.11; MCH: 0.74 ± 0.11; ACV: 0.79 ± 0.05; Shock: 0.85 ± 0.09; Figure 6E). Together, these results support the view that the compartmental pattern of cAMP dynamics is broadly reflected in downstream PKA activity, consistent with the functional relevance of the cAMP code.

**Figure 6.**
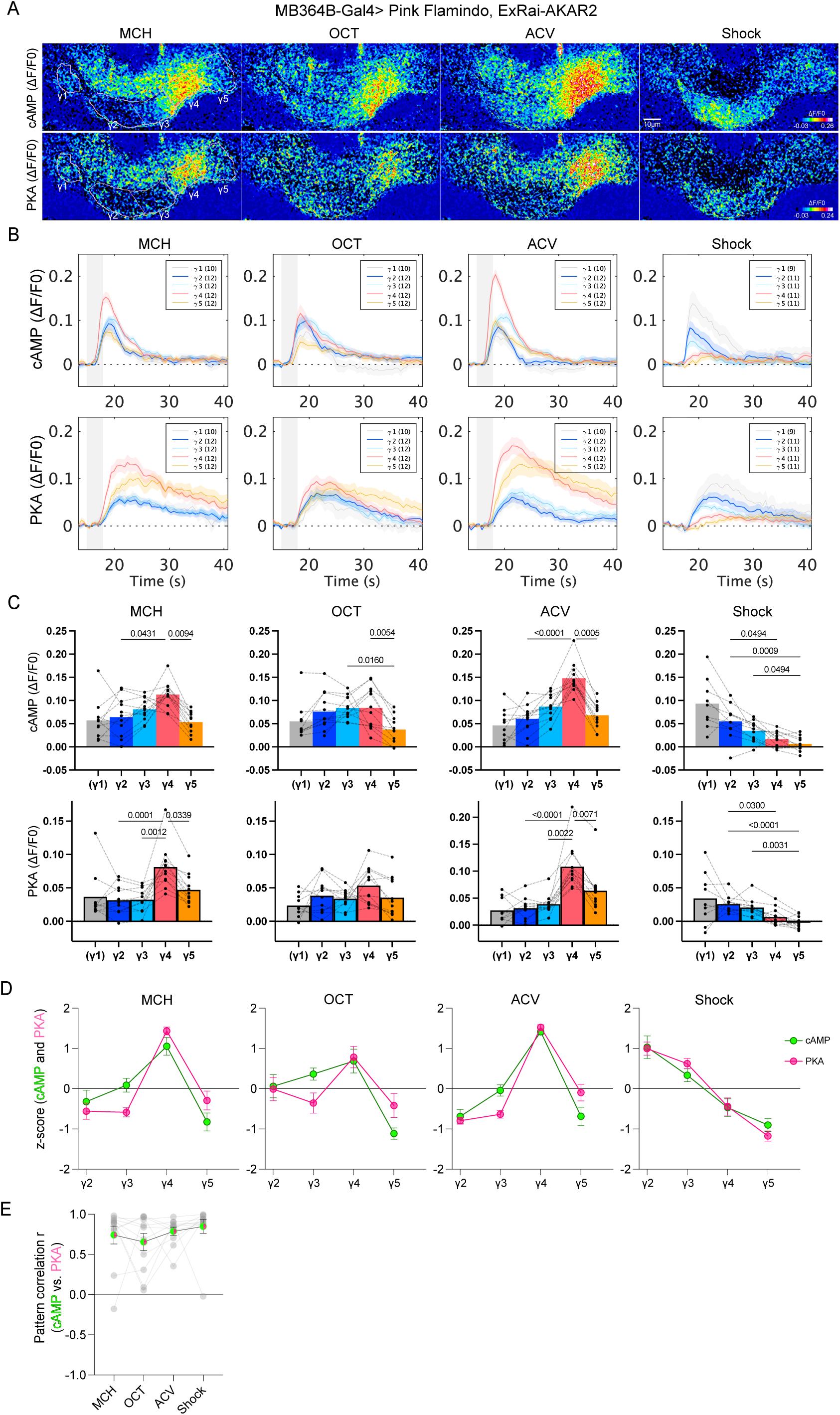
PKA activity broadly reflects the compartmental cAMP pattern. (A) Averaged ΔF/F0 images from flies expressing Pink Flamindo for cAMP detection and ExRai-AKAR2 for PKA activity detection under the control of the MB364B-GAL4 driver are shown. cAMP and PKA signals were simultaneously recorded. ROIs for each compartment are overlaid on the left panels. Mean peak ΔF/F0 images averaged over 5 s (10 frames) in the γ1–γ5 compartments during OCT, MCH, ACV, and shock presentation are also shown. (B) A series of line plots depicting the temporal dynamics of cAMP (upper panels) and PKA activity (lower panels) over time in seconds (s) across different stimulus conditions. Each subplot corresponds to a specific stimulus and experimental stage. The y-axis indicates ΔF/F0. The solid lines represent the mean values for each compartment, calculated from the number of flies indicated in parentheses within the subplots. The shaded areas around each line represent the SEM. Shaded rectangles indicate the estimated period of odor delivery to the fly (approximately 16–19 s). A 60-V electric shock was delivered at 17.5 s for 10 ms. (C) Quantification of averaged peak ΔF/F0 images (5 s, 10 frames) of cAMP (upper panels) and PKA activity (lower panels). Data from different compartments derived from the same animal are connected by dashed lines. Bar graphs represent mean values. Values from the γ1 compartment are shown in parentheses for reference but were not included in the statistical analyses. Responses across the γ2–γ5 compartments were compared using the Friedman test followed by Dunn’s multiple comparisons test for all pairwise comparisons. p-values are shown only for comparisons with significant differences. For MCH, OCT, and ACV, γ1, n = 10 and γ2–γ5, n = 12. For shock, γ1, n = 9 and γ2–γ5, n = 11. (**D**) Within-sample z-scored cAMP and PKA profiles (γ2–γ5) for each of four stimuli — OCT, MCH, ACV and electric shock presented alone — overlaid on the same axes for each stimulus (n = 11; only flies with responses to all four stimuli were included). Mean ± SEM per stimulus. (**E**) Pattern correlation r between cAMP and PKA for each of the four stimuli. Small circles show individual flies (n = 11); colored circles and error bars show mean ± SEM. Mean r exceeded 0.6 for every stimulus (OCT: 0.65 ± 0.11; MCH: 0.74 ± 0.11; ACV: 0.79 ± 0.05; Shock: 0.85 ± 0.09). Figure 6A-video (related to Figure 6A-C). Figure 6B-video (related to Figure 6B and C). Videos illustrating simultaneous cAMP and PKA imaging, acquired at 2 Hz and played back at 6 Hz.

### The Rutabaga adenylyl cyclase mutant *rut¹* lacks detectable conditioning-associated reorganization and shows an altered pre-conditioning cAMP code

The Rutabaga adenylyl cyclase has been regarded as the coincidence detector, as described above, and has indeed been demonstrated to increase cAMP levels synergistically only in the presence of both the CS and US (Gervasi et al., 2010; Tomchik and Davis, 2009). We failed to observe the anticipated synergistic elevation of cAMP levels upon pairing CS and US in flies with wild-type *rutabaga*; instead, we observed an increase that was at most additive (Figure 3B). Nevertheless, given the pivotal role of rut in aversive memory formation, we still contemplated what *rut* mutants might reveal.

The *rutabaga^1^* (*rut^1^*) allele, which is well known for its distinctive learning deficit phenotype in the context of aversive memory, was initially isolated from the X chromosome collection mutagenized by ethyl-methanesulfonate (Livingstone et al., 1984). This allele carries a point mutation that results in loss of detectable Rutabaga adenylyl cyclase activity (Levin et al., 1992). We monitored cAMP transients using G-Flamp1 in the *rut^1^*flies during aversive conditioning, following the protocol described above, with OCT serving as the conditioned stimulus (Figure 7B, D, F, J and L). Across the individual odor-response profiles, cAMP responses in *rut¹* flies were not globally reduced relative to those in the genetic control (Figure 7A–F), consistent with previous reports that Rutabaga accounts for only a portion of cAMP production in KCs (Noyes and Davis, 2023; Tomchik and Davis, 2009). In the genetic control, OCT-evoked cAMP responses after conditioning were smaller in the γ4 compartment, whereas those in γ2 were not reduced (Figure 7E). During conditioning, OCT-evoked cAMP responses in control flies increased in γ2 and decreased in γ4 relative to Pre, and cAMP responses to ACV (CS−) in γ2 also increased (Figure 7G); none of these changes reached significance in *rut¹* flies (Figure 7H). Accordingly, the γ2/γ4 cAMP ratio increased significantly from Pre to Post for OCT (p = 0.0156), but not for MCH (p = 0.3828) or ACV (p = 0.3125; Figure 7I). In *rut¹* flies, by contrast, the γ2/γ4 cAMP ratio did not change significantly for any odor (MCH, p = 0.2500; OCT, p = 0.9102; ACV, p = 0.2031; Figure 7J). To visualize these changes at the level of the compartmental profile, we compared the within-sample z-scored cAMP profile (γ1–γ5) across the phases of conditioning between the genetic control and *rut¹* flies (Figure 7K, L). In control flies, conditioning altered the relative distribution of OCT (CS+)-evoked cAMP across compartments: relative cAMP levels increased in γ2 and decreased in γ4 (Figure 7K). In *rut¹* flies, by contrast, the OCT profile remained largely unchanged, with similar Pre and Post values in both compartments (Figure 7L). Why was this conditioning-associated reorganization not observed in *rut¹* flies? To address this question, we compared pre-conditioning responses between genotypes. Because the genotype-dependent differences were broadly consistent in direction across MCH, OCT, and ACV, responses to the three odors were averaged within each fly. In this analysis, the γ2/γ4 cAMP ratio was higher in *rut¹* flies than in the genetic control (p = 0.0184; Figure 7M). Consistent with this, mean odor-evoked cAMP responses in *rut¹* flies were significantly lower in the γ4 compartment (p = 0.0474), whereas those in γ2 were numerically higher but not significantly different (p = 0.3031; Figure 7N). Thus, in *rut¹* flies, the compartmental balance of odor-evoked cAMP responses already differed from that of controls before conditioning. Although mean shock-evoked cAMP responses in the γ4 compartment were significantly higher in *rut¹* flies than in the genetic control (p = 0.0078; Figure 7O), this difference was not reproduced with the alternative cAMP sensor Pink Flamindo (Figure 8X). We therefore did not pursue this observation further. Whether the absence of conditioning-associated reorganization in *rut¹* mutants reflects a failure to reorganize or an already shifted starting state cannot be distinguished from these data. Either way, the lack of reorganization is in line with the well-established impairment of aversive memory formation in *rut¹* mutants.

**Figure 7.**
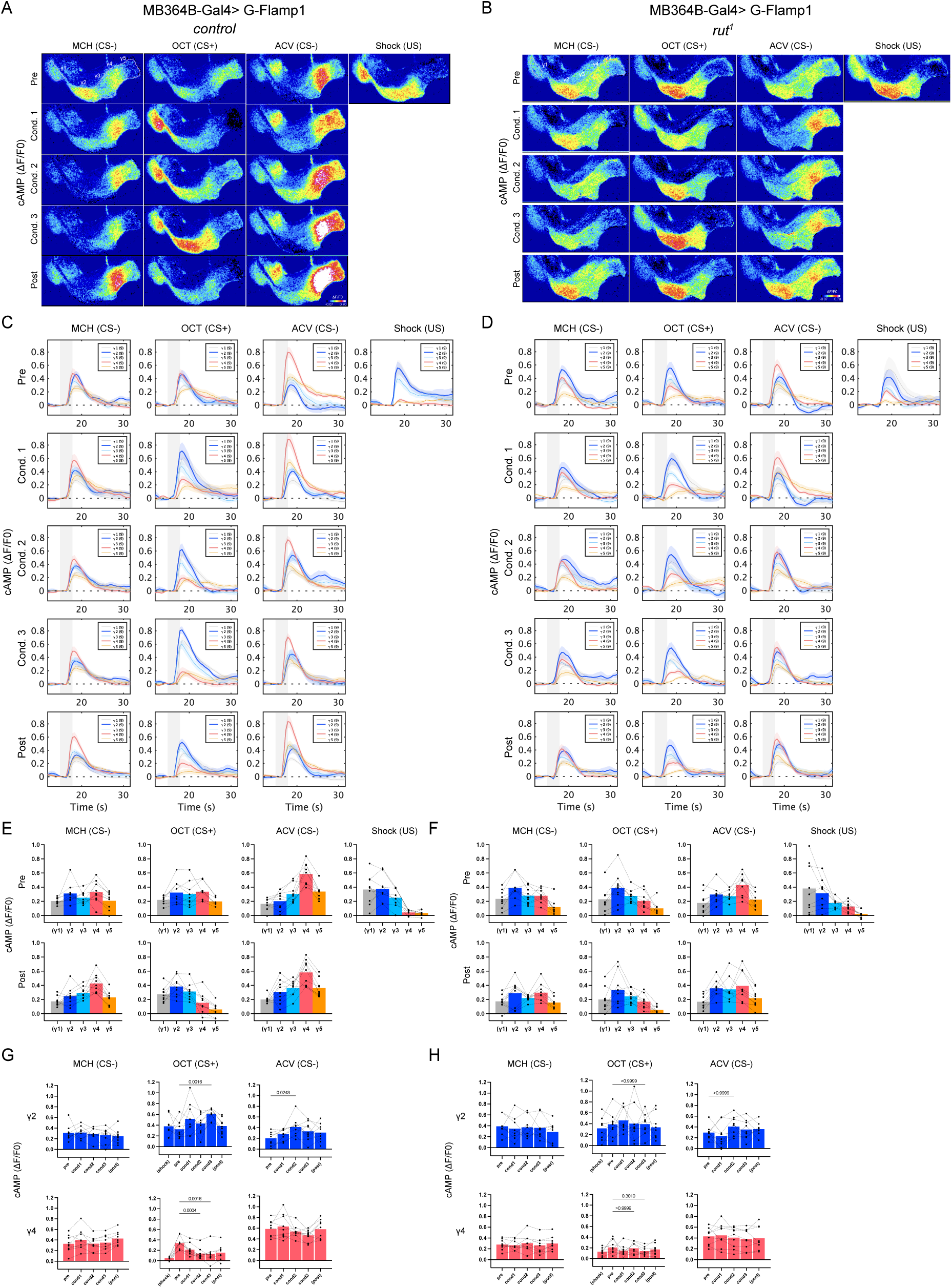
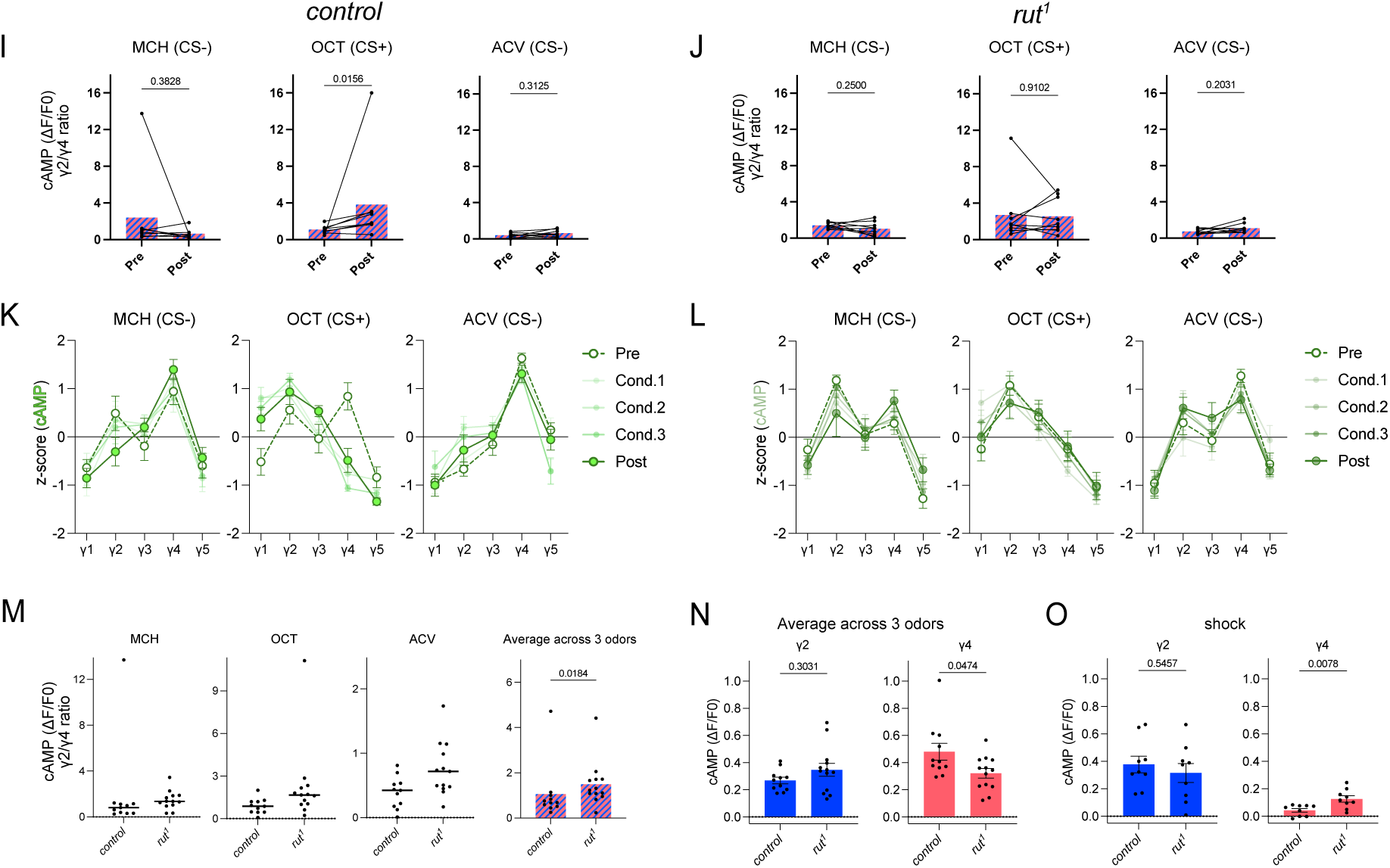
The Rutabaga adenylyl cyclase mutant *rut¹* lacks detectable conditioning-associated reorganization and shows an altered pre-conditioning cAMP code. (**A, B**) The ΔF/F0 images, showing the averaged peak responses (10 frames over 5 s) of cAMP in the γ1–γ5 compartments of KCs, were captured during the presentations of MCH (CS−), OCT (CS+), ACV (CS−), and shock (US) in a genetic control fly **(A**) and a *rut¹* mutant fly (B) throughout conditioning. The ROIs for each compartment are mapped onto the top left panel. (**C, D**) Temporal dynamics of cAMP in the γ1–γ5 compartments for each stimulus and phase in control (C) and *rut¹* (D) flies. The solid lines represent the mean ΔF/F0 for each compartment, calculated from the number of flies indicated in parentheses within the subplots, and the shaded areas represent the SEM. Shaded rectangles indicate the estimated period of odor delivery to the fly (approximately 16–19 s). A 60-V electric shock was delivered at 17.5 s for 10 ms. (**E, F**) Quantification of the averaged ΔF/F0 images of control (E) and *rut¹* (F) flies. The plots of different compartments derived from the same sample are connected with dashed lines. The bar graphs represent mean values. n = 9 (E) and 9 (F). (**G, H**) Changes in cAMP levels in the γ2 and γ4 compartments across conditioning steps in control (G) and *rut¹* (H) flies. cAMP responses were compared using the Friedman test followed by Dunn’s multiple comparisons test, with the corresponding pre-conditioning odor response used as the reference condition. The same datasets used in (E) and (F) were used for (G) and (H), respectively. Shock-evoked responses in the CS+ experimental group, as well as post-conditioning responses to each stimulus, are shown for reference but were not included in the statistical analyses. In G, p-values are shown for comparisons with significant differences. In H, p-values are shown for the comparisons that were significant in G; none of the comparisons in *rut¹* flies reached significance. n = 9 for all stimulus sets in both γ2 and γ4. (**I, J**) The ratio of cAMP levels in the γ2 compartment to those in the γ4 compartment was calculated for each odor response in control (I) and *rut¹* (J) flies. Bar graphs represent mean values. Ratios before and after conditioning were compared using the Wilcoxon signed-rank test, and p-values are shown for all comparisons. The same datasets used in (E) and (F) were used for (I) and (J), respectively. One control fly was excluded from the γ2/γ4 cAMP ratio analysis because its γ4 response to post-conditioning OCT (CS+) was negative. n = 8 (I) and 9 (J). (K, L) Within-sample z-scored cAMP profiles (γ1–γ5) for MCH (CS−), OCT (CS+), and ACV (CS−) across Pre, Cond.1–3, and Post in control (K) and *rut¹* (L) flies (n = 9 per genotype). Because γ1 responses were obtained from all flies in this dataset, γ1 was included in these profiles. Mean ± SEM at each phase. Corresponding phase legends are shown alongside each panel. (M) γ2/γ4 ratio of pre-conditioning odor-evoked cAMP responses in control and *rut¹* flies, shown for each odor and averaged across the three odors within each fly. Each data point represents one fly; horizontal lines indicate the median, and the bar graph represents the mean. Data for individual odors are shown for reference; statistical testing was performed on responses averaged across the three odors. p-value, Mann–Whitney U test; control, n = 11; *rut¹*, n = 13. (N) Comparison of pre-conditioning odor-evoked cAMP responses in the γ2 and γ4 compartments between control and *rut¹* flies. Responses to MCH, OCT, and ACV were averaged within each fly before statistical comparison. Each data point represents the mean response to the three odors from one fly. Bar graphs represent mean values ± SEM. p-values are shown. Mann–Whitney U test; control, n = 11; *rut¹*, n = 13. Data in (M) and (N) include flies subjected to the conditioning paradigm (pre-conditioning responses) and additional flies in which only odor responses were recorded without conditioning. (O) Comparison of shock-evoked cAMP responses in the γ2 and γ4 compartments between control and *rut¹* flies. Shock-evoked responses were recorded in the shock-alone trial. Each data point represents one fly. Bar graphs represent mean values ± SEM. p-values are shown. Mann–Whitney U test; control, n = 9; *rut¹*, n = 9. Figure 7A-video (related to Figure 7A, C, E, G, I and K). Figure 7B-video (related to Figure 7B, D, F, H, J and L). Figure 7C-video (related to Figure 7C, E, G, I and K). Figure 7D-video (related to Figure 7D, F, H, J and L). Videos illustrating cAMP imaging, acquired at 2 Hz and played back at 6 Hz.

**Figure 8.**
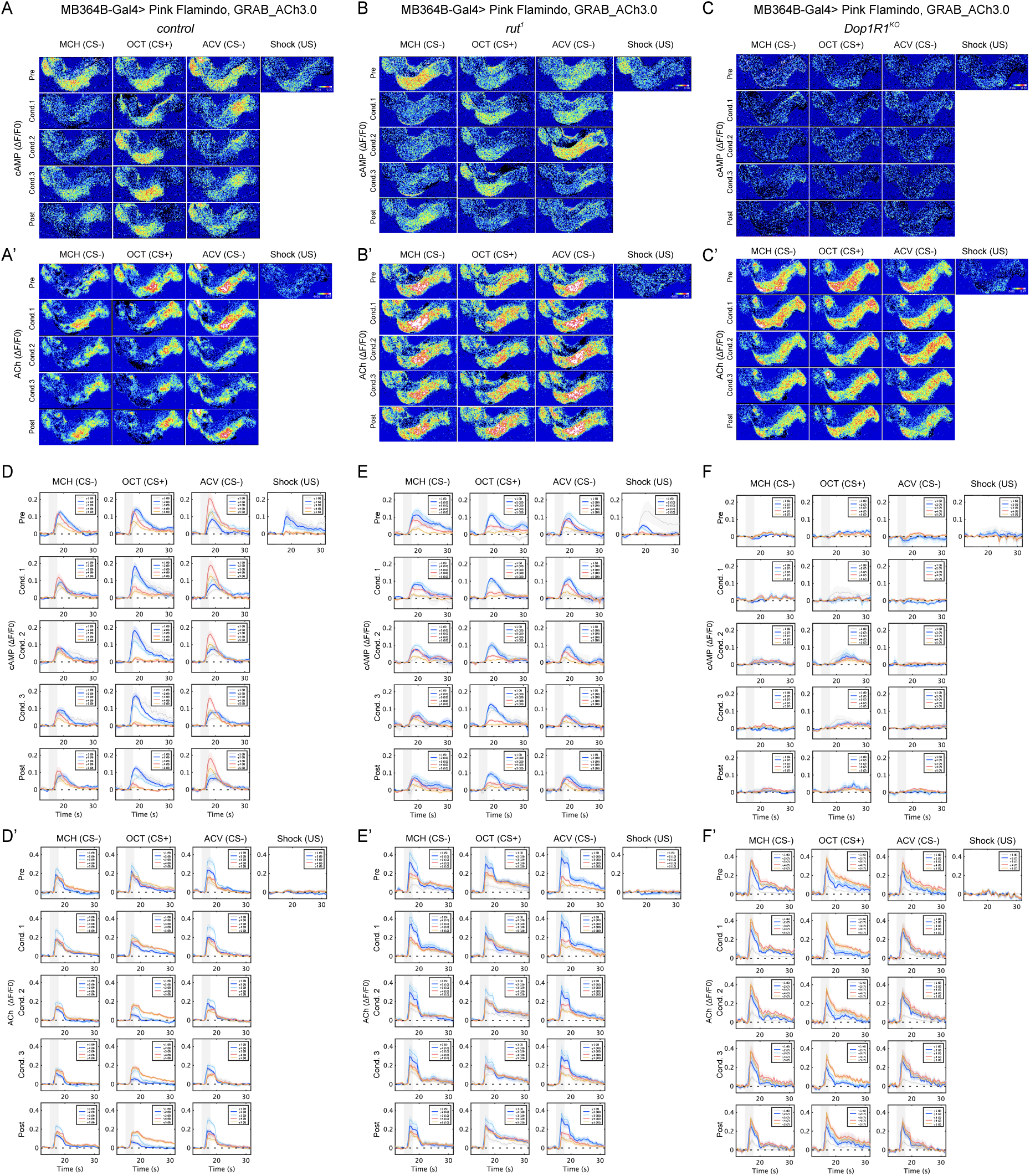
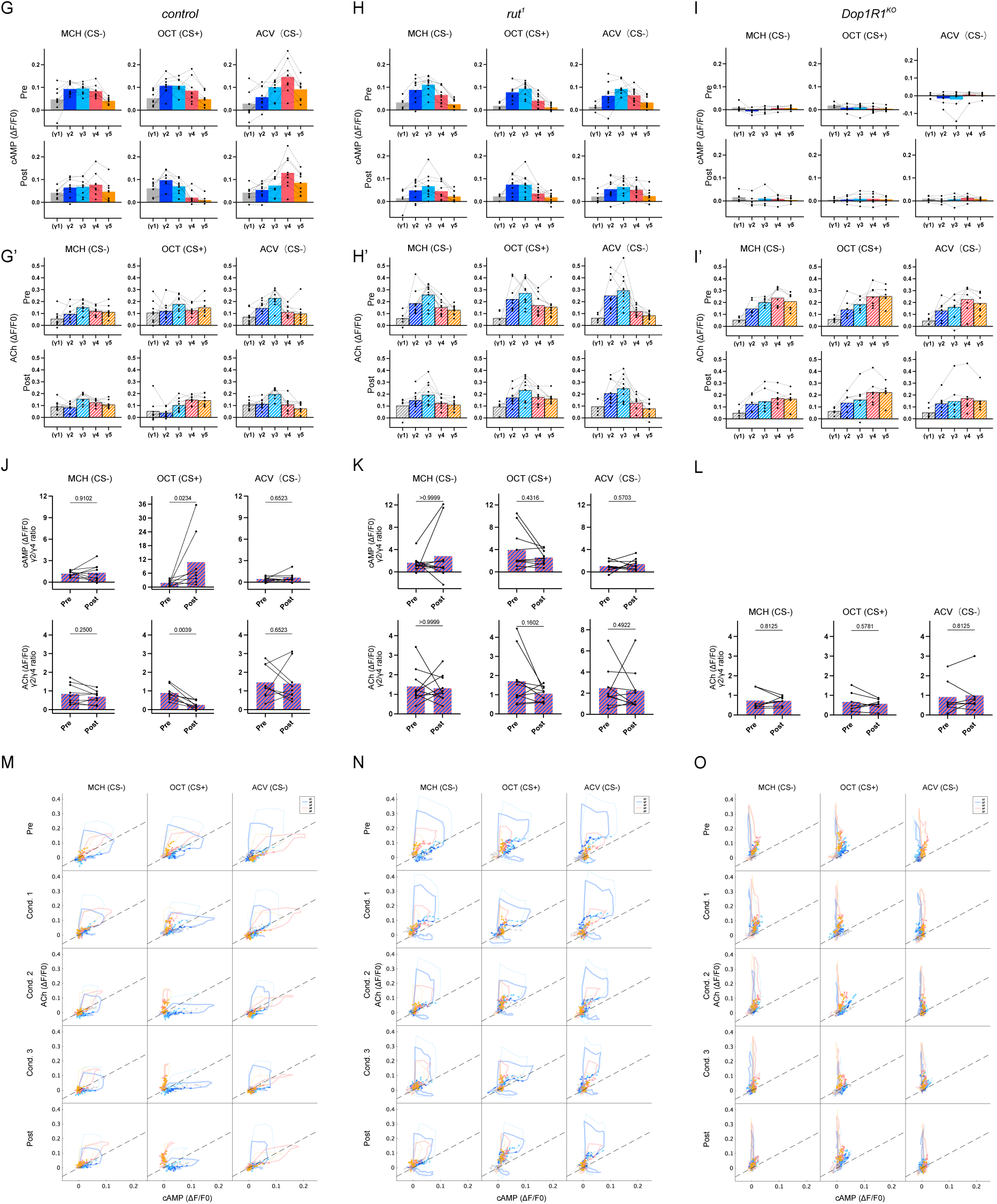
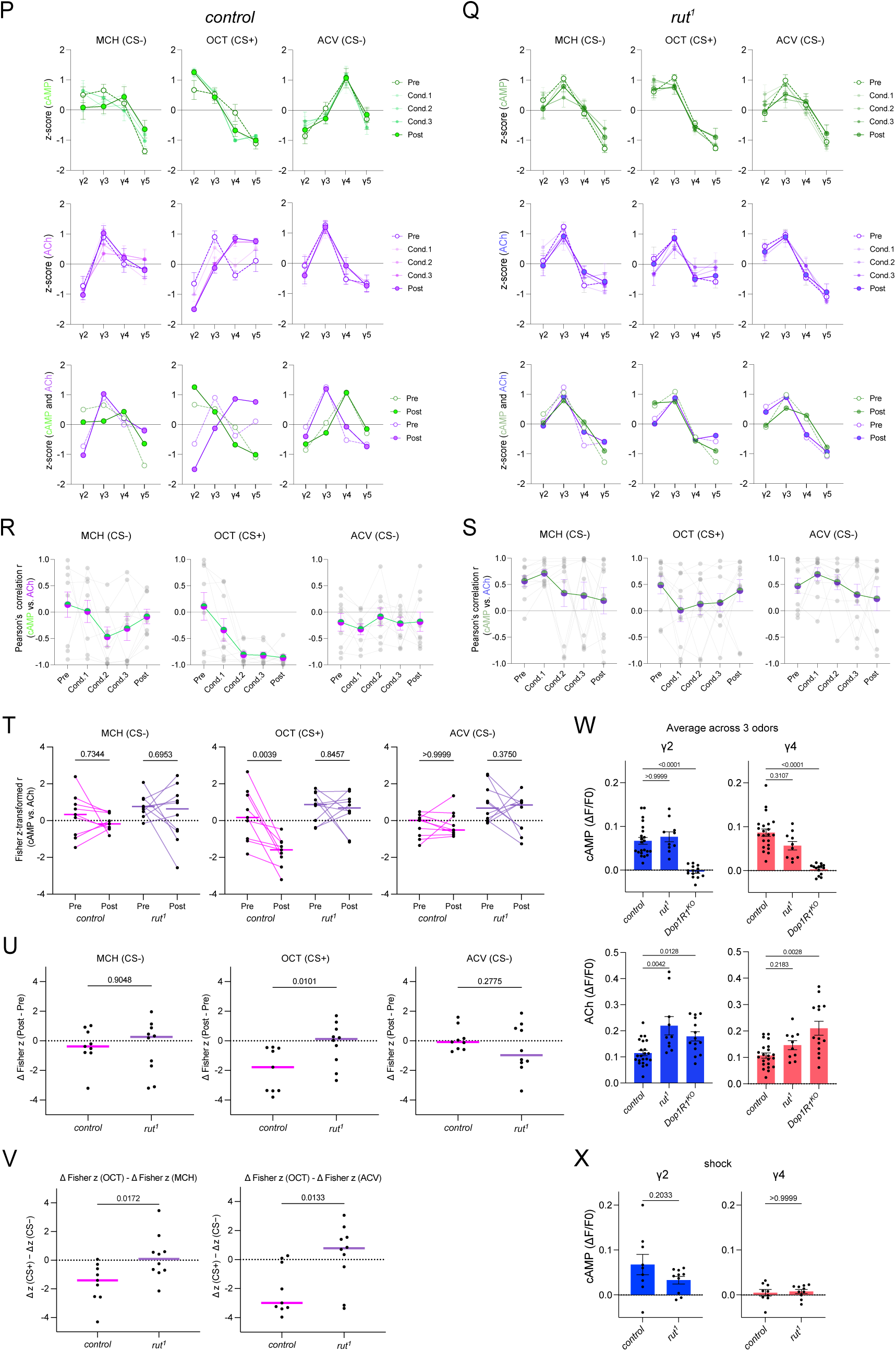
Aversive conditioning induces an inverse relationship between the compartmental cAMP code and ACh release for the CS+ in control but not in *rut¹* flies. All cAMP and ACh images in the γ1–γ5 compartments of KCs were simultaneously captured from flies expressing Pink Flamindo and GRAB_ACh3.0 under the control of the MB364B-GAL4 driver and subjected to aversive conditioning, following the procedure outlined in Figure 4. (A-C, A′-C′) The ΔF/F0 images, depicting the averaged peak responses (10 frames over 5 s) of cAMP (A–C) and ACh (A′-C′) in the γ1–γ5 compartments of KCs, were captured during the presentations of MCH (CS−), OCT (CS+), ACV (CS−), and shock (US) throughout conditioning in control (A, A′), *rut¹* (B, B′), and *Dop1R1^KO^* (C, C′) flies. The ROIs for each compartment are mapped onto the top left panel. (D-F, D′-F′) Temporal dynamics of simultaneously recorded cAMP (D-F) and ACh (D′-F′) in the γ1–γ5 compartments for each stimulus and phase in control (D, D′), *rut¹* (E, E′), and *Dop1R1^KO^* (F, F′) flies. The solid lines represent the mean ΔF/F0 for each compartment, calculated from the number of flies indicated in parentheses within the subplots, and the shaded areas represent the SEM. Shaded rectangles indicate the estimated period of odor delivery to the fly (approximately 16–19 s). A 60-V electric shock was delivered at 17.5 s for 10 ms. (G–I, G′–I′) Quantification of the averaged peak ΔF/F0 images of pre- and post-conditioning responses to odors for cAMP (G–I) and ACh (G′–I′) in control (G, G′), *rut¹* (H, H′), and *Dop1R1^KO^* (I, I′) flies. The plots of different compartments derived from the same sample are connected with dashed lines. The bar graphs represent mean values. n = 9 (G, G′), 10 (H, H′), and 7 (I, I′). (J–L) γ2/γ4 ratios calculated from the datasets in (G, G′), (H, H′), and (I, I′), respectively. Upper panels, cAMP; lower panels, ACh. Ratios before and after conditioning were compared using the Wilcoxon signed-rank test, and p-values are shown for all comparisons. The γ2/γ4 cAMP ratio was not calculated for *Dop1R1^KO^* flies because the γ4 response was negative in several trials, precluding meaningful interpretation of the ratio (see I). cAMP: n = 9, 8, 9 (J) and 10, 10, 9 (K) for MCH, OCT, and ACV, respectively. ACh: n = 9, 9, 9 (J), 10, 10, 10 (K), and 7, 7, 7 (L). For each odor, flies in which the γ4 cAMP response was negative in either condition were excluded from the ratio analysis. (M–O) Trajectories of cAMP (x-axis) and ACh (y-axis) ΔF/F0 values over time for the γ1–γ5 compartments in control (M), *rut¹* (N), and *Dop1R1^KO^* (O) flies. Each point along a trajectory represents the mean cAMP and ACh responses across flies at each time point, calculated from the datasets in (D, D′), (E, E′), and (F, F′), respectively. The γ2 and γ4 compartments are highlighted with thicker lines, and arrowheads indicate the direction of time. The dashed line (y = x) is shown as a visual reference. (P, Q) Within-sample z-scored profiles (γ2–γ5) of cAMP (top) and ACh (middle) for MCH (CS−), OCT (CS+), and ACV (CS−) across Pre, Cond.1–3, and Post in control (P) and *rut¹* (Q) flies. Mean ± SEM at each phase; n = 9 for control and 10 for *rut¹*. Corresponding phase legends are shown alongside each panel. Bottom panels: the Pre and Post profiles of cAMP and ACh are overlaid on the same axes. *Dop1R1^KO^* flies were not included in the analyses in (P–V) because their cAMP responses were near zero. (R, S) Pattern correlation r between cAMP and ACh profiles across phases for MCH (CS−), OCT (CS+), and ACV (CS−) in control (R) and *rut¹* (S) flies. Small grey circles show individual flies (n = 9 for control and 10 for *rut¹*); colored circles and error bars show mean ± SEM. (T) Fisher z-transformed pattern correlation between cAMP and ACh at Pre and Post for each odor in control and *rut¹* flies. Lines connect data from the same fly (magenta, control; purple, *rut¹*); horizontal bars indicate the median. p-values are shown for all comparisons (two-sided Wilcoxon signed-rank test). n = 9 (control) and 10 (*rut¹).* (U) Change in the Fisher z-transformed pattern correlation from Pre to Post (Δz = Post − Pre) for each odor, compared between control and rut¹ flies. Each data point represents one fly; horizontal bars indicate the median. p-values are shown for all comparisons (Mann–Whitney U test). n = 9 (control) and 10 (*rut¹*). (V) Difference between the change for OCT and that for each CS− odor (Δz(OCT) − Δz(CS−)), compared between control and rut¹ flies. Each data point represents one fly; horizontal bars indicate the median. p-values are shown for both comparisons (Mann–Whitney U test). n = 9 (control) and 10 (*rut¹*). This contrast served as the primary test of odor selectivity; the comparisons in (T) and (U) are shown for descriptive purposes. (W) Comparison of pre-conditioning odor-evoked cAMP (top) and ACh (bottom) responses in the γ2 and γ4 compartments among control, *rut¹*, and *Dop1R1^KO^* flies. Responses to MCH, OCT, and ACV were averaged within each fly before statistical comparison. Each data point represents the mean response to the three odors from one fly. Bar graphs represent mean values ± SEM. Kruskal–Wallis test followed by Dunn’s multiple comparisons test against control; Dunn-adjusted p-values are shown, and values below 0.0001 are reported as p < 0.0001. Control, n = 22; *rut¹*, n = 10; *Dop1R1^KO^*, n = 14. For control and *Dop1R1^KO^*, data include flies subjected to the conditioning paradigm (pre-conditioning responses) and additional flies in which only odor responses were recorded without conditioning. (X) Comparison of shock-evoked cAMP responses in the γ2 and γ4 compartments between control and *rut¹* flies, measured with Pink Flamindo. Shock-evoked responses were recorded in the shock-alone trial. Each data point represents one fly. Bar graphs represent mean values ± SEM. p-values are shown. Mann–Whitney U test; Control, n = 9; *rut¹*, n = 10. Figure 8AA′-video (related to Figure 8A, A′, D, D′, G, G′, J, M, P, R and T–W). Figure 8BB′-video (related to Figure 8B, B′, E, E′, H, H′, K, N, Q, S and T–W). Figure 8CC′-video (related to Figure 8C, C′, F, F′, I, I′, L, O and W). Figure 8DD′-video (related to Figure 8D, D′, G, G′, J, M, P, R and T–W). Figure 8EE′-video (related to Figure 8E, E′, H, H′, K, N, Q, S and T–W). Figure 8FF′-video (related to Figure 8F, F′, I, I′, L, O and W). Videos illustrating simultaneous cAMP and acetylcholine imaging, acquired at 2 Hz and played back at 6 Hz.

### Aversive conditioning induces an inverse relationship between the compartmental cAMP code and ACh release for the CS+ in control but not in *rut¹* flies

The KCs are recognized as cholinergic (Barnstedt et al., 2016; Yasuyama and Salvaterra, 1999), and their ACh release after aversive conditioning is reportedly attenuated in the γ1 (Zeng et al., 2023), γ2 and γ3 (Stahl et al., 2022; Zeng et al., 2023), and α′3 compartments (Zhang et al., 2019). To examine the relationship between cAMP dynamics and ACh release, we conducted simultaneous imaging of ACh and cAMP transients in the γ KCs of flies expressing the ACh sensor GRAB_ACh3.0 (Jing et al., 2020) and Pink Flamindo under the control of the MB364B-GAL4 driver, while the flies were subjected to aversive conditioning under the microscope (Figure 8). As observed with G-Flamp1 (Figure 2C and F), cAMP responses were nearly abolished in *Dop1R1*^KO^ mutants, whereas ACh responses were preserved (Figure 8C, C′, F, F′, I and I′), and the γ2/γ4 ACh ratio did not change significantly for any odor (Figure 8L; see also Figure 8O). In the genetic control, after conditioning, cAMP responses to the CS+ odor OCT were markedly reduced in the γ4 and γ5 compartments (Figure 8A, D and G), as observed with G-Flamp1 (Figure 7E). Accordingly, the γ2/γ4 cAMP ratio increased for OCT (p = 0.0234; Figure 8J, top), replicating the result obtained with G-Flamp1 (Figure 7I) with an independent sensor. ACh responses to OCT, in contrast, decreased within the γ1 and γ2 compartments but remained constant within the γ4 and γ5 compartments (Figure 8A′, D′ and G′), leading to a decrease in the γ2/γ4 ACh ratio (p = 0.0039; Figure 8J, bottom). Neither ratio changed significantly for the CS− odors MCH and ACV (cAMP: p = 0.9102 and 0.6523; ACh: p = 0.2500 and 0.6523, respectively; Figure 8J). When cAMP and ACh responses were plotted together in a two-dimensional plot (Figure 8M), relative ACh levels did not exhibit a uniform inverse relationship with cAMP levels across compartments and conditions. After conditioning, however, responses to the CS+ odor showed an increase in the γ2/γ4 cAMP ratio together with a decrease in the γ2/γ4 ACh ratio, revealing a conditioning-dependent relationship between the compartmental cAMP pattern and ACh release from KCs. To examine this relationship across compartments, we plotted the within-sample z-scored compartmental profiles (γ2–γ5) of cAMP and ACh across the phases of conditioning (Figure 8P). *Dop1R1^KO^*mutants were not included in this and the following correlation analyses, because their cAMP responses were near zero. In control flies, the cAMP and ACh profiles for OCT (CS+) diverged in shape after conditioning: cAMP increased in γ2, where ACh decreased, whereas relative ACh levels increased in γ4 and γ5 (Figure 8P, top and middle). This inverse relationship became unambiguous when the two profiles were directly overlaid (Figure 8P, bottom). For most other odor–phase combinations, the two signals showed no clear relationship by visual inspection alone. We quantified this relationship with the pattern correlation r between the cAMP and ACh profiles. In control flies, r for OCT decreased progressively during conditioning, becoming strongly negative from Cond.2 onward and remaining so through Post (Figure 8R). In contrast, r for the CS− odors showed no sustained change. In individual flies, the Fisher z-transformed r for OCT decreased from Pre to Post in every control fly (p = 0.0039), whereas no change was observed for MCH (p = 0.7344) or ACV (p > 0.9999; Figure 8T).

In the *rut¹* background (Figure 8B, B′, E, E′, H, H′, K and N), neither the cAMP nor the ACh γ2/γ4 ratio changed significantly for any odor after conditioning (cAMP: MCH, p > 0.9999; OCT, p = 0.4316; ACV, p = 0.5703; ACh: MCH, p > 0.9999; OCT, p = 0.1602; ACV, p = 0.4922; Figure 8K). The response trajectories of cAMP and ACh in *rut¹* flies were notably uniform across odors, regardless of the conditioning phase (Figure 8N), in contrast to the diverse response patterns observed in the genetic control (Figure 8M). The inverse relationship seen in control flies was absent in *rut¹* flies; the cAMP and ACh profiles remained similar at both Pre and Post (Figure 8Q). Accordingly, r for OCT did not show the sustained negative shift seen in controls: it decreased at Cond.1, remained low through Cond.3, and partially recovered by Post (Figure 8S). The Fisher z-transformed r did not change from Pre to Post for any odor in rut¹ flies (MCH, p = 0.6953; OCT, p = 0.8457; ACV, p = 0.3750; Figure 8T).

To determine whether the conditioning-induced change in pattern correlation differed between genotypes, we directly compared control and *rut¹* flies. The change in pattern correlation from Pre to Post (Δz = Post − Pre) differed between genotypes only for OCT (p = 0.0101), not for MCH (p = 0.9048) or ACV (p = 0.2775; Figure 8U). To test odor selectivity, we compared the difference between the change for OCT and that for each CS− odor (Δz (OCT) − Δz(CS−)) between genotypes. This difference was significantly more negative in control than in *rut¹* flies for both CS− odors (OCT − MCH, p = 0.0172; OCT − ACV, p = 0.0133; Figure 8V). Thus, the decrease in cAMP–ACh pattern correlation occurred selectively for the CS+ relative to both CS− odors and was absent in *rut¹* flies.

To examine how Rutabaga and Dop1R1-dependent dopaminergic signaling affect KC cholinergic output, we also compared pre-conditioning odor-evoked cAMP and ACh responses among the genetic control, *rut¹*, and *Dop1R1^KO^* mutant flies (Figure 8W). As in Figure 7, responses to the three odors were averaged within each fly. In *rut¹* flies, cAMP responses in the γ4 compartment were numerically lower than in the genetic control, as observed with G-Flamp1 (Figure 7N), but the difference did not reach significance (p = 0.3107); no difference was observed in γ2 (p > 0.9999). No significant differences in shock-evoked cAMP responses were observed between *rut¹* flies and the genetic control (Figure 8X). ACh responses in *rut¹* flies were significantly higher in γ2 (p = 0.0042), but not in γ4 (p = 0.2183). In *Dop1R1*^KO^ mutants, cAMP responses were nearly abolished in both compartments (both p < 0.0001), whereas ACh responses were significantly higher in both γ2 (p = 0.0128) and γ4 (p = 0.0028). The increase in ACh responses in *Dop1R1* mutants differs from a previous report of reduced ACh levels under *Dop1R1* knockdown conditions (Noyes and Davis, 2023). Thus, loss of either Rutabaga or Dop1R1 was associated with larger odor-evoked ACh responses before conditioning, although the compartments affected differed between the two mutants. Notably, in *rut¹* flies, ACh responses were increased in the γ2 compartment, where cAMP responses did not differ significantly from those in the genetic control, suggesting that Rutabaga may influence ACh release through mechanisms not fully captured by bulk cAMP levels.

### Elevating cAMP suppresses ACh release from KCs

The larger ACh responses in *Dop1R1^KO^* mutants are consistent with negative regulation of KC cholinergic output by cAMP signaling, although the enhanced odor-evoked Ca^2+^ responses in these mutants (Figure 2H) may also contribute. To test directly whether elevating cAMP is sufficient to suppress ACh release, we monitored ACh release while activating adenylyl cyclases by bath application of forskolin (Figure 9). Upon forskolin application, cAMP levels increased markedly, whereas ACh levels decreased in the γ2 compartment (Figure 9A, B); no such changes were observed with vehicle (Figure 9B, left). Forskolin also reduced odor-evoked ACh responses (Figure 9C). For OCT, responses in the γ2–γ5 compartments were significantly reduced (γ2, p = 0.0119; γ3, p = 0.0054; γ4, p = 0.0119; γ5, p = 0.0009) and recovered after washout (Pre vs Washout: γ2–γ4, p > 0.9999; γ5, p = 0.6346; Figure 9D). Together, these results indicate that elevating cAMP can suppress KC cholinergic output and that aversive conditioning reorganizes the relationship between the compartmental cAMP pattern and ACh release.

**Figure 9.**
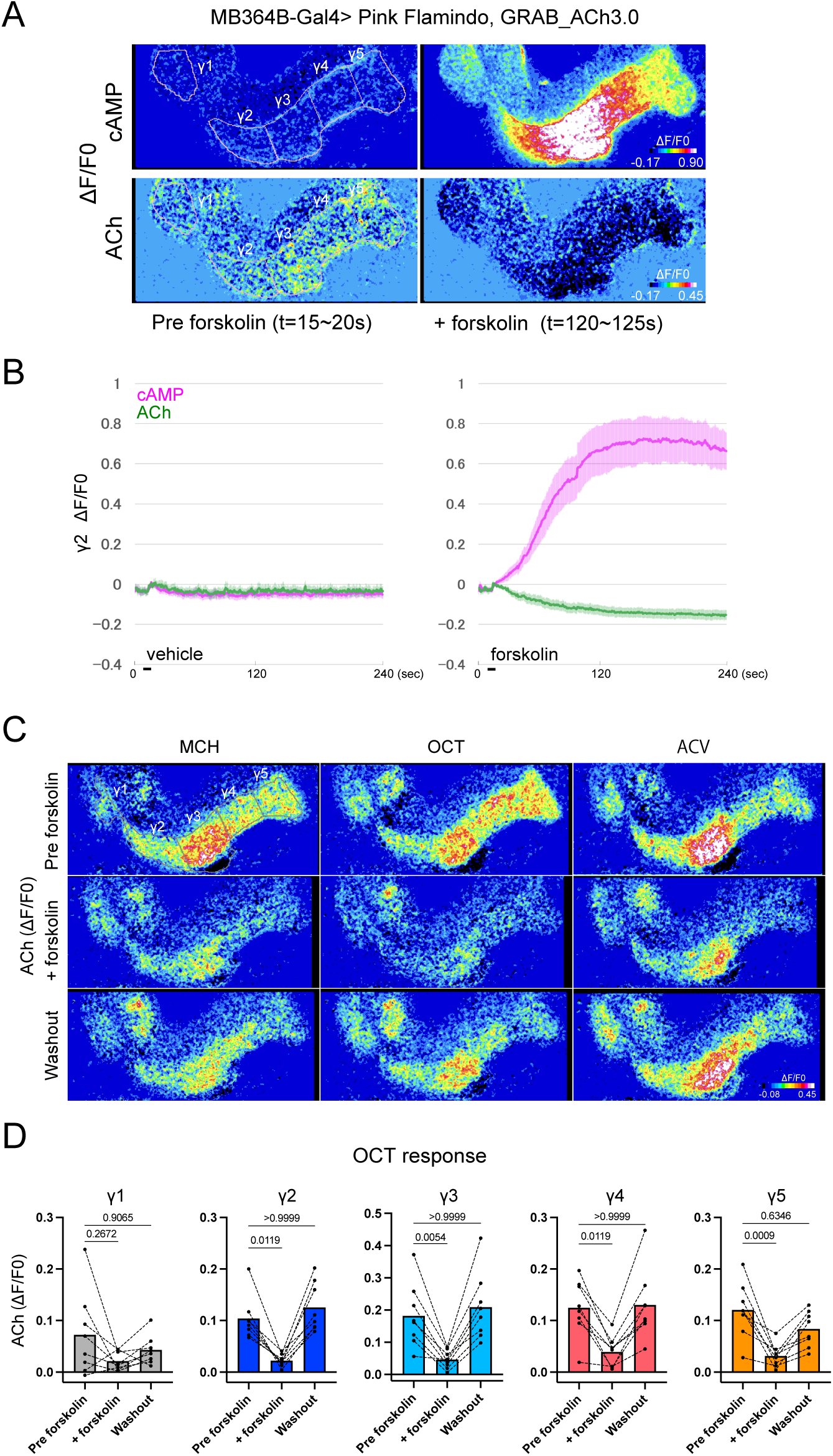
Elevating cAMP suppresses ACh release from KCs. To examine the effect of adenylyl cyclase activation on ACh release, cAMP and ACh signals were monitored during bath application of forskolin in flies expressing Pink Flamindo and GRAB_ACh3.0 under the control of the MB364B-GAL4 driver. (A) The ΔF/F0 images of cAMP and ACh were obtained by averaging the responses over 5 s (10 frames) immediately before switching from vehicle (saline containing 0.5% EtOH) to saline containing 100 μM forskolin and 0.5% EtOH (Pre forskolin; left panels) and 2 min after the switch (forskolin; right panels). The ROIs for each compartment are mapped onto the left panels. (B) Time-series traces of cAMP (magenta) and ACh (green) in the γ2 compartment show the changes in their levels after switching from standard saline to vehicle solution containing 0.5% EtOH (left) or from vehicle to saline containing 100 μM forskolin and 0.5% EtOH (right). The short black lines indicate the periods of solution exchange; thereafter, the preparation was maintained in the indicated vehicle- or forskolin-containing saline for the remainder of the recording. Solid lines and shaded areas represent the mean ± SEM. n = 7 for vehicle and n = 8 for forskolin. (C) ΔF/F0 images of averaged peak ACh responses over 5 s (10 frames) during the presentation of OCT, MCH, or ACV before forskolin application (Pre forskolin), 4 min after switching from vehicle to saline containing 100 μM forskolin (forskolin), and following the return to standard saline (Washout). The ROIs for each compartment are mapped onto the top left panel. (D) The averaged peak ΔF/F0 responses of ACh to OCT under the pre-forskolin, forskolin, and washout conditions, as depicted in (C), were quantified for each compartment. The plots of different conditions derived from the same sample are connected with dashed lines. The bar graph illustrates the mean values. The three conditions were compared using the Friedman test followed by Dunn’s multiple comparisons test, with the Pre forskolin response used as the reference condition. p-values are shown for all comparisons. γ1-γ5; n = 8. Data in Figure 9A–D were obtained from the same cohort of eight flies. For the vehicle condition in Figure 9B, one recording was excluded because substantial brain movement during acquisition precluded reliable fluorescence quantification; therefore, n = 7 for vehicle and n = 8 for forskolin. Figure 9A-video (related to Figure 9A and B). Figure 9B-video (related to Figure 9B). Videos illustrating ACh and cAMP imaging, acquired at 2 Hz and played back at 6 Hz. Figure 9C-video (related to Figure 9C and D). Figure 9D-video (related to Figure 9D). Videos illustrating ACh imaging, acquired at 2 Hz and played back at 6 Hz.

## Discussion

Our results indicate that, at the resolution of our imaging, cAMP levels do not by themselves distinguish CS+ responsive KCs from the remaining KC population. Instead, our data suggest that cAMP relays dopaminergic reinforcement signals to KC output: elevating cAMP suppresses ACh release from KCs, and conditioning establishes an inverse relationship between the compartmental cAMP code and ACh release for the CS+. Through this action, cAMP may modulate the DAN-KC-MBON recurrent feedback loop (Adel and Griffith, 2021; Cognigni et al., 2018; Felsenberg et al., 2018, 2017; Jovanoski et al., 2023; Kato et al., 2023; Li et al., 2020; Modi et al., 2020; Otto et al., 2020; Rubin and Aso, 2024) in a valence-dependent manner. We propose that the cAMP levels across the γ compartments of the mushroom body constitute a “cAMP code,” representing a KC-intrinsic, compartmentalized expression of integrated valence. Rather than encoding CS+ identity at the single-KC level, this cAMP code serves to link dopaminergic input to changes in KC cholinergic output within the DAN-KC-MBON circuit, thereby contributing to the formation or modulation of memory engrams.

### Bulk cAMP levels do not distinguish CS+ responsive KCs from other KCs

Physiological and cellular studies on Aplysia gill withdrawal (Abrams and Kandel, 1988; Ocorr et al., 1985; Yovell et al., 1992) and genetic studies on Drosophila olfactory conditioning (Heisenberg, 2003; Levin et al., 1992; Livingstone et al., 1984; Mons et al., 1999; Tully, 1987; Tully and Quinn, 1985) solidified the idea of adenylyl cyclase as a coincidence detector. A similar framework has been proposed in vertebrates: CS-evoked calcium and US-evoked dopamine converge on calcium-sensitive adenylyl cyclase to drive reward learning in the striatum and nucleus accumbens (Urakubo et al., 2020; Yagishita et al., 2014; Yamaguchi et al., 2022). Consistent with this, calcium-sensitive adenylyl cyclase is necessary for mossy-fiber LTP (Villacres et al., 1998; Wang et al., 2003) and for dopamine-dependent LTP in CA1 (Fuchsberger et al., 2025).

To directly evaluate this long-standing hypothesis, we quantified in vivo cAMP with concurrent calcium or dopamine transients at near–single-cell resolution during Drosophila aversive olfactory memory formation and observed inconsistencies. Odor (CS) evokes dopamine release from overlapping DAN populations that also mediate shock (US) signals. Consistent with this, we did not observe a selective increase in cAMP levels in CS+ responsive KCs, even in the γ1 and γ2 compartments, which are necessary and sufficient for the acquisition of short- and middle-term memories (Aso et al., 2012). These results challenge the notion that cAMP levels alone are sufficient to define a cellular engram for the Drosophila olfactory conditioning (Figure 10A).

**Figure 10.**
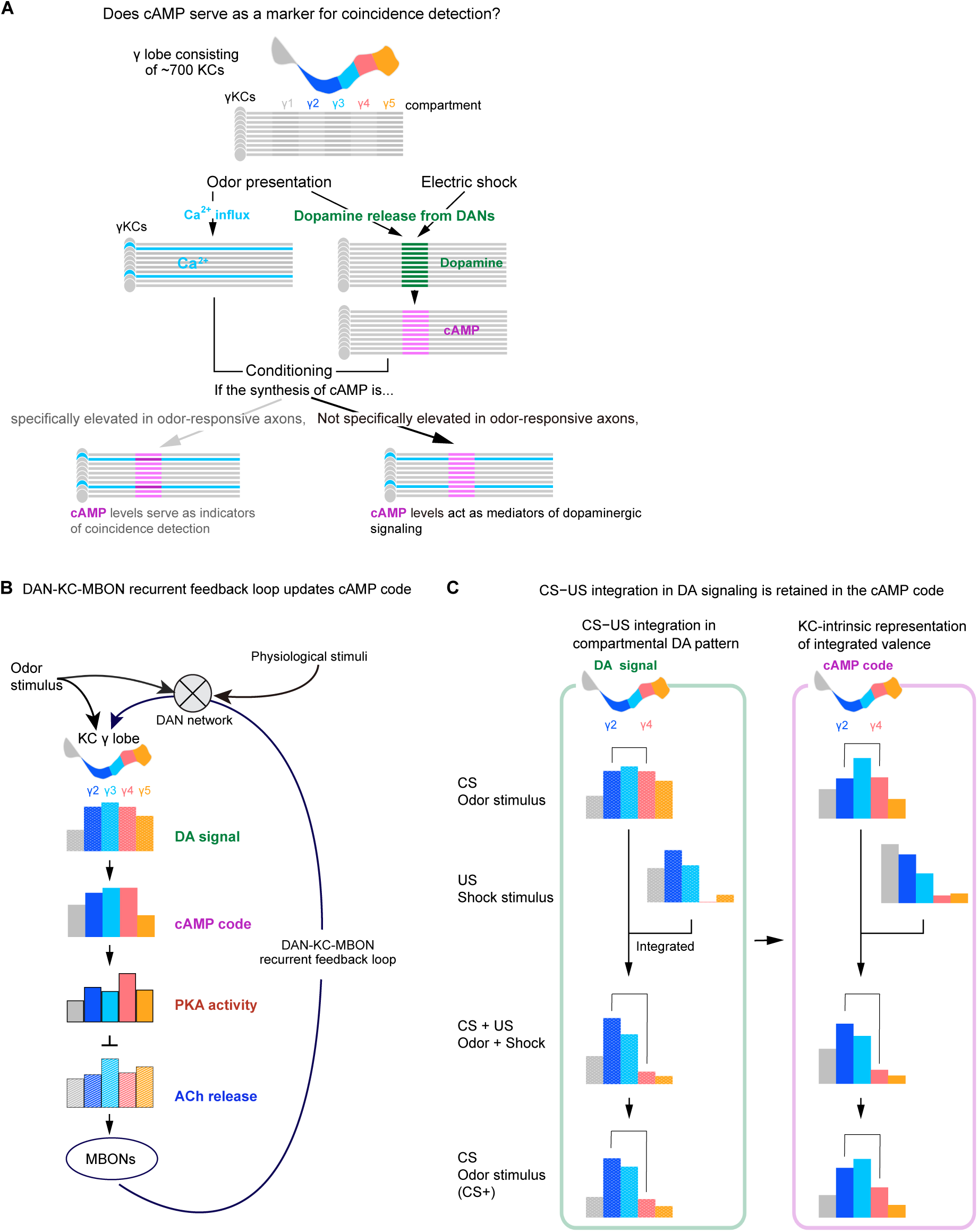
A model for cAMP signaling in olfactory memory formation. (A) Does cAMP serve as a marker for coincidence detection? The γ lobe comprises approximately 700 Kenyon cells (KCs) and is partitioned into five compartments, and each receives its own distinct dopaminergic innervation. Each olfactory stimulus generally elicits calcium transients in a unique subset of approximately 5–9% of KCs (Honegger et al., 2011) and activates a specific subset of dopaminergic neurons. Consequently, KCs receive compartment-specific dopamine signals along the entire length of the lobe, exhibiting distinct characteristics for each odor. The release of dopamine occurs through volume transmission, leading to its detection in a population of axons that spans nearly the entire breadth of the lobe. For simplicity, only the dopamine signal within the γ2 compartment is depicted. This broadly distributed dopamine signal approximately coincides spatially with the cAMP signal. If the synthesis of cAMP is significantly enhanced by calcium, then the axons of odor-responsive KCs with high calcium signals exhibit elevated cAMP synthesis. Conversely, if calcium does not predominantly regulate cAMP synthesis, then the axons activated by olfactory stimuli are not distinguished by their cAMP levels. Our findings support the latter hypothesis and suggest that cAMP serves as a mediator of the dopamine signal. (B) The DAN-KC-MBON recurrent feedback loop updates the cAMP code. cAMP levels in each compartment depend on dopaminergic input conveying negative valence to the γ1-3 compartments and positive valence to the γ4-5 compartments. The global equilibrium of cAMP levels throughout the γ lobe could yield a comprehensive depiction of the integrated valence, which can be portrayed as a two-dimensional image of cAMP levels and designated as the cAMP code. KCs exert regulatory control over the functions of their downstream targets, namely the mushroom body output neurons (MBONs), through the release of acetylcholine (ACh). cAMP negatively regulates ACh release, possibly through PKA activity. In turn, the MBONs reciprocally modulate dopaminergic activities, thereby establishing a complex DAN-KC-MBON recurrent feedback loop. The DAN network also receives inputs reflecting physiological states and environmental cues. The OCT-evoked responses are derived from Figures 5 (dopamine), 6 (cAMP and PKA) and 8 (ACh). (C) CS– US integration is reflected in compartmental dopamine signaling. When the odor is paired with electric shock, relative dopamine responses increase in the γ2 compartment and decrease in the γ4 compartment, a pattern that is subsequently reflected in the cAMP code. This suggests that the conditioned stimulus (odor) and unconditioned stimulus (electric shock) are integrated at the level of dopaminergic signaling. The responses to OCT and shock presentations are derived from Figure 5.

### Alternative models for coincidence detection

The Rutabaga AC is not necessarily involved in all phases of aversive olfactory memory, as it is crucial for labile anesthesia-sensitive memory (ASM), but dispensable for anesthesia-resistant memory (ARM). In contrast, the phosphodiesterase Dunce is essential for ARM, but not for ASM, despite both Rutabaga and Dunce serving in the cAMP pathway (Scheunemann et al., 2012). In addition, Rutabaga is required concurrently in both α/β and γ KCs, whereas Dunce functions in both KCs and antennal lobe local neurons (Scheunemann et al., 2012). Another olfactory memory paradigm, known as "trace conditioning," does not require the Rutabaga AC and allows the presentation of the conditioned stimulus (CS) and the unconditioned stimulus (US) to be temporally separated by intervals of up to 60 s (Shuai et al., 2011). The Rutabaga AC has been identified as the sole type-I AC in Drosophila (Livingstone, 1985), and its activation by calcium and the Gαs subunit has led to its postulation as a coincidence detector for the CS and US. This implies that the underlying mechanism for the formation of ARM and trace conditioning relies on another AC or an alternative mechanism. Nair et al. presented a theoretical framework wherein the cAMP/PKA signal activated by dopamine converges with the calcium signal at CaMKII, when the inputs of calcium and dopamine are delivered in temporal proximity and in a dopamine-following-calcium sequence (Nair et al., 2016). This role of CaMKII could underlie the case of Drosophila olfactory conditioning, since CaMKII accumulation in the KC axons is required for aversive and appetitive conditioning (Adel et al., 2022; Chen et al., 2022; but see also Yamada et al., 2024). Yamada et al. recently demonstrated that the targeted application of forskolin alone does not induce presynaptic plasticity (Yamada et al., 2024), manifested as long-term depression (LTD), which is characteristically observed at the KC-MBON synapse after the associative pairing of olfactory cues with aversive stimuli (Hige et al., 2015). However, when the optogenetic stimulation of KCs is coupled with the application of forskolin, LTD is indeed induced, implying that these two inputs converge somewhere downstream of cAMP synthesis (Yamada et al., 2024). They also found no evidence for synergistic action of dopamine and optogenetic KC activation on cAMP synthesis, consistent with our observation that cAMP did not increase supra-additively upon CS–US pairing (Figure 3B). The forskolin-induced reduction of ACh release observed here recovered after washout (Figure 9D), consistent with an acute modulation of release rather than LTD.

Several studies have suggested alternative mechanisms of coincidence detection. Saitoe et al. proposed that the aversive input of electric shocks is conveyed by glutamate, but not dopamine; rather, dopamine release is elicited after this pairing event and serves solely as a reinforcement, but not the US (Saitoe, 2021; Ueno et al., 2017). They further demonstrated that the glutamate transmission is mediated by the ensheathing glia (EG) that surround the MBs (Miyashita et al., 2023). Qiao et al. demonstrated that the PPL1 DANs modulate the input-timing-dependent plasticity of γ KC synapses from projection neurons, depending on type 2 dopamine receptor (DD2R) (Qiao et al., 2022). Their findings further indicated that the activation of the PPL1-DD2R pathway elicits a rise in calcium levels, which is dependent on the voltage-gated calcium channel (VGCC). They concluded that the VGCC could serve as the coincidence detector. However, this conclusion does not extend to all phases of aversive memory, as the authors also demonstrated that DD2R is not necessary for short-term memory formation.

The potential role of the DAN-KC-MBON recurrent feedback loop in the formation of olfactory memory raises the prospect that the synaptic connections between KCs and MBONs may also rely on spike-timing-dependent plasticity (STDP) (Zeng et al., 2023). In the locust, Cassenaer and Laurent showed that pre–post pairing that induces STDP at KC– MBON synapses can, when combined with local application of the reinforcement-signaling neuromodulator octopamine, tag those synapses for associative modification (Cassenaer and Laurent, 2012). This exemplifies a three-factor (neo-Hebbian) learning rule. In this rule, the coincidence of pre- and postsynaptic activity marks synapses as eligible for change, and a third, neuromodulatory signal decides whether that change is expressed (Gerstner et al., 2018). In Drosophila, dopamine is the most likely third factor. DANs convey reinforcement signals to the mushroom body, and connectomic analyses have revealed direct synaptic contacts from DANs onto MBONs in the α lobe (Takemura et al., 2017). Pribbenow et al. have indeed demonstrated that concurrent dopamine release and focal acetylcholine application to M6 (MBON-γ5β′2a) dendrites, circumventing KC activity, induce plasticity in calcium transients within this MBON (Pribbenow et al., 2022). Zeng et al. further found that the dorsal paired medial (DPM) neuron releases 5-HT on KCs and regulates the coincidence time window, which has been hypothesized by several theories to be a timing regulator (Zeng et al., 2023).

### Dopaminergic population dynamics may underlie aversive olfactory memory formation

Upon exposure to different odors, distinct populations of DANs exhibit specific activity patterns, as documented in multiple studies (Berry et al., 2012; Huang et al., 2024; Kato et al., 2023; Mao and Davis, 2009; Yamagata et al., 2016). This, in turn, may shape the cAMP code in the γ lobe, which may contribute to the conditioning-associated suppression of ACh release (Figure 10B). Given that the sustained population dynamics among DANs, likely facilitated by the interactions within the DAN-KC-MBON recurrent feedback loop, integrate changes in physiological states and environmental cues (Huang et al., 2024; Kato et al., 2023; Siju et al., 2020), associative memory formation may be understood as one component of this broader dynamical process. This is consistent with the additional finding that repeated exposure to an aversive odor can give rise to a labile, self-reinforced aversive memory for that odor (Jacob et al., 2021). If both odor and electric shock stimuli are represented within the cAMP-code framework, electric shock may be viewed as an extreme aversive signal within that framework, since both olfactory stimuli (CS) and electric shock (US) activate overlapping populations of DANs. The imaging of dopamine levels further supports this view. During conditioning, the compartmental pattern of dopamine evoked by OCT paired with shock closely resembled that evoked by shock alone, as expected, and the concurrently recorded cAMP pattern followed it (Figure 5K). After conditioning, the similarity of the OCT-evoked cAMP pattern to the shock-evoked dopamine and cAMP patterns remained significantly higher than before conditioning, and the corresponding increase for dopamine showed a similar trend (Figure 5L). These results suggest that the CS and US converge in compartmental dopamine signaling during conditioning, and that this convergence is retained in the cAMP code after conditioning (Figure 10C). In other words, the cAMP code for the CS+ odor is shifted toward the shock-evoked pattern relative to its pre-conditioning state, suggesting that pairing integrates this code with that of the shock US, which carries a much more negative valence. Assuming this to be true, the question of where and how the learned population activity of DANs is stored may be closely related to the broader question of where and how odor-specific population activity patterns of DANs are represented and maintained.

In addition to the well-established function of the exogenously activated γ1-pedc DAN as the aversive US (Aso et al., 2012; Aso and Rubin, 2016), the suppression of the corresponding MBON, γ1-pedc>α/β, can also serve as an aversive US (König et al., 2019; Ueoka et al., 2017), as noted (Yamada et al., 2023). These discoveries suggest that not only the stimulation of the γ1pedc DAN, but also the inhibition of γ1pedc>α/β MBON can trigger a series of events that tilt DAN population activity towards an aversively learned state.

Assuming that the population activities of DANs are continuously shaped by the DAN-KC-MBON recurrent feedback loop that integrates inputs from relevant compartments, alterations in the γ1 compartment resulting from increased γ1 DAN activity or decreased γ1 MBON activity could trigger a large-scale reshaping of DAN population activity.

Despite the current lack of knowledge regarding the higher-order mechanisms responsible for the organization of the DAN-KC-MBON recurrent feedback loop, recent progress in unraveling the hierarchical architecture of the DAN circuit (Rubin and Aso, 2024; Yamada et al., 2023) holds promise for illuminating these mechanisms in the near future.

Although CS and US signals may both be represented in compartmental patterns of dopamine signaling, their integration during associative learning is unlikely to reflect a simple summation of these signals, because associative learning depends critically on the relative input timing of CS and US. Thus, identifying the molecular basis of coincidence detection is essential for fully understanding the mechanisms underlying associative learning.

### The role of Rutabaga AC

The *rut^1^* mutation has been shown to abolish Rutabaga AC activity (Levin et al., 1992). However, in the *rut^1^* mutant, cAMP levels were found to be either comparable to or slightly lower than those in the control strain. This finding suggests that substantial cAMP levels in KCs are maintained by other adenylyl cyclases. It has indeed been shown that innate odor preferences depend on another adenylyl cyclase, ACXD (Noyes and Davis, 2023). Nonetheless, short-term aversive memory is impaired in the *rut^1^*mutant, and this deficit can be ameliorated by expressing wild-type Rutabaga in γ and α/β KCs (McGuire et al., 2003; Zars et al., 2000). This underscores the crucial role of Rutabaga in the formation of aversive memory within γ and α/β KCs.

In our current imaging analyses of γ KCs, the altered innate odor-evoked cAMP code in *rut¹* provides a potential link to the olfactory learning defect of *rut¹*. The compartmental configuration of odor-evoked cAMP responses is already altered even before conditioning. Notably, the pre-conditioning γ2/γ4 ratio in *rut¹* was shifted in the same direction as the increase in this ratio associated with conditioning-induced plasticity in control flies. This altered starting configuration may reduce the dynamic range available for further γ2/γ4 reorganization during conditioning, thereby contributing to the absence of a significant Pre-to-Post change observed in *rut¹*. This scenario would arise if Rutabaga’s contribution to cAMP synthesis is not uniform across compartments in wild-type flies. Under this view, the compartments differ in how heavily they normally rely on Rutabaga for their baseline cAMP levels, so that removing Rutabaga does not simply scale cAMP down uniformly across the mushroom body; instead, it disproportionately reduces cAMP in the compartments most dependent on it, thereby reconfiguring the relative (compartmental) balance of the cAMP code — and potentially changing which compartment dominates the baseline configuration. We currently do not know whether Rutabaga also plays an additional, instructive role in conditioning-dependent plasticity beyond setting this baseline configuration. Finally, whether the Ca^2+^ sensitivity of Rutabaga is required for its physiological function remains an open question that cannot be resolved by the present results.

**Shock-paired conditioning enhances the DA–cAMP pattern correlation for the CS+**

Pattern correlation analysis revealed that conditioning had a substantial effect on the correlation between cAMP and DA. Before conditioning, odor-evoked DA and cAMP profiles were moderately correlated, with considerable variation between individuals, and the correlation was lowest for OCT (Figure 5I). During conditioning, by contrast, this correlation for OCT rose markedly, and remained elevated when OCT was presented alone after conditioning (Post, Figure 5J), reaching a level comparable to that for the CS− odors. Although the high correlation during conditioning may partly reflect the strong shock-evoked DA signal, its persistence at Post, in the absence of shock, cannot be explained in this way. Why does this correlation increase? One possibility is that a threshold exists, above which cAMP more readily tracks DA levels; alternatively, a non-linear mechanism may be at work. Beyond a purely quantitative relationship between DA level and cAMP synthesis, we cannot rule out the involvement of an as-yet-unidentified physiological mechanism—for example, a coincidence-detection process—that becomes engaged specifically under these conditions.

### An inverse cAMP–ACh relationship emerges with CS+ conditioning

Because elevating cAMP suppresses ACh release (Figure 9), cAMP and ACh might be expected to show an inverse compartmental relationship regardless of conditioning history. Before conditioning, however, such a relationship was not evident: with odor presentation alone, many individuals showed positive pattern correlations, and values varied considerably across flies. With CS+ conditioning, by contrast, a clear inverse relationship emerged and was maintained through the post-conditioning period, suggesting that it may constitute a component of the memory trace. For OCT, this trajectory forms a mirror image of the DA– cAMP pattern correlation described above. These observations suggest that the compartmental mirror-image relationship reported here reflects a conditioning-induced state rather than a constitutive, always-on property of cAMP–ACh coupling. Whether this phenomenon depends solely on cAMP levels or involves additional mechanisms, such as a requirement for coincident KC activity downstream of cAMP synthesis (Yamada et al., 2024), remains unknown. Unlike control flies, *rut¹* mutants did not develop the conditioning-induced inverse cAMP–ACh correlation, and this genotype difference was significant (Figure 8U, V). However, because *rut¹* flies already show an altered cAMP code before conditioning and lack its conditioning-associated reorganization (Figure 7), it remains unclear from this observation alone whether Rutabaga is required for the coupling between cAMP and ACh release itself, and further study will be needed.

### Limitations of the study

In this study, our conclusions regarding the correlation between cAMP and calcium were drawn solely from live imaging, which was constrained by the temporal and spatial resolution of the microscope and affected by sample movement during imaging. Although the close correspondence between the compartmental patterns of PKA activity and cAMP suggests that the detected cAMP signals were accompanied by PKA activation, we cannot rule out that the principal locus of Rutabaga-dependent coincidence detection occurs in an inaccessible microdomain, or that Rutabaga-derived cAMP at the coincidence site was masked by cAMP produced by other adenylyl cyclases. Further refinement of imaging techniques, along with complementary approaches such as analyses using mutants, may help to more precisely elucidate the correlation between the two signals. Because CS− odors were always presented after the shock-paired CS+ trials and an explicitly unpaired control group was not included, we cannot fully exclude non-associative contributions, such as sensitization or adaptation, to the changes observed during conditioning. However, the persistent shift at Post was specific to the paired odor regardless of odor identity and was absent in PPL1-silenced and *rut¹* flies subjected to the identical protocol. In addition, although the forskolin-induced decrease in ACh release was reversible and was not observed with vehicle, we cannot fully exclude cAMP-independent actions of forskolin or its actions on cells other than KCs, as forskolin was bath-applied and an inactive analog was not tested.

## Materials and methods

### Fly strains

Fly strains were maintained on standard food containing cornmeal at 25°C. The following strains were used: γCRE-p-GAL4 i.e., CRE-p65-AD/CyO; MB247-GAL4-DBD (Yamazaki et al., 2018), UAS-G-Flamp1 attP40/CyO and UAS-G-Flamp1 VK00005 (gifts from Jun Chu and Yulong Li, Nat. Commun., 2022.13:5363), TH-LexA VK00040 (a gift from Yoshinori Aso), UAS-jRGECO1b VK00005 (BDSC #63795), MB364B-GAL4 i.e., R13F02-p65-AD attP40; R21B06-GAL4-DBD attP2 (BDSC #68318), OK107-GAL4 (#Kyoto DGRC 106098), UAS-Pink Flamindo su(Hw) attP5 and UAS-Pink Flamindo VK00027 (this study), UAS-GRAB_DA2m attP40 (BDSC #90878), *Dop1R1^KO^* (D1R1 KO, a gift from Taro Ueno), UAS-ExRai-AKAR2 VK00005 (this study), UAS-GRAB_ACh3.0 attP40 (BDSC #86549), *rut¹* (BDSC #9404), and LexAop-TNT (a gift from Chi-Hon Lee; Karuppudurai et al., 2014).

### Fly genotypes used by figures

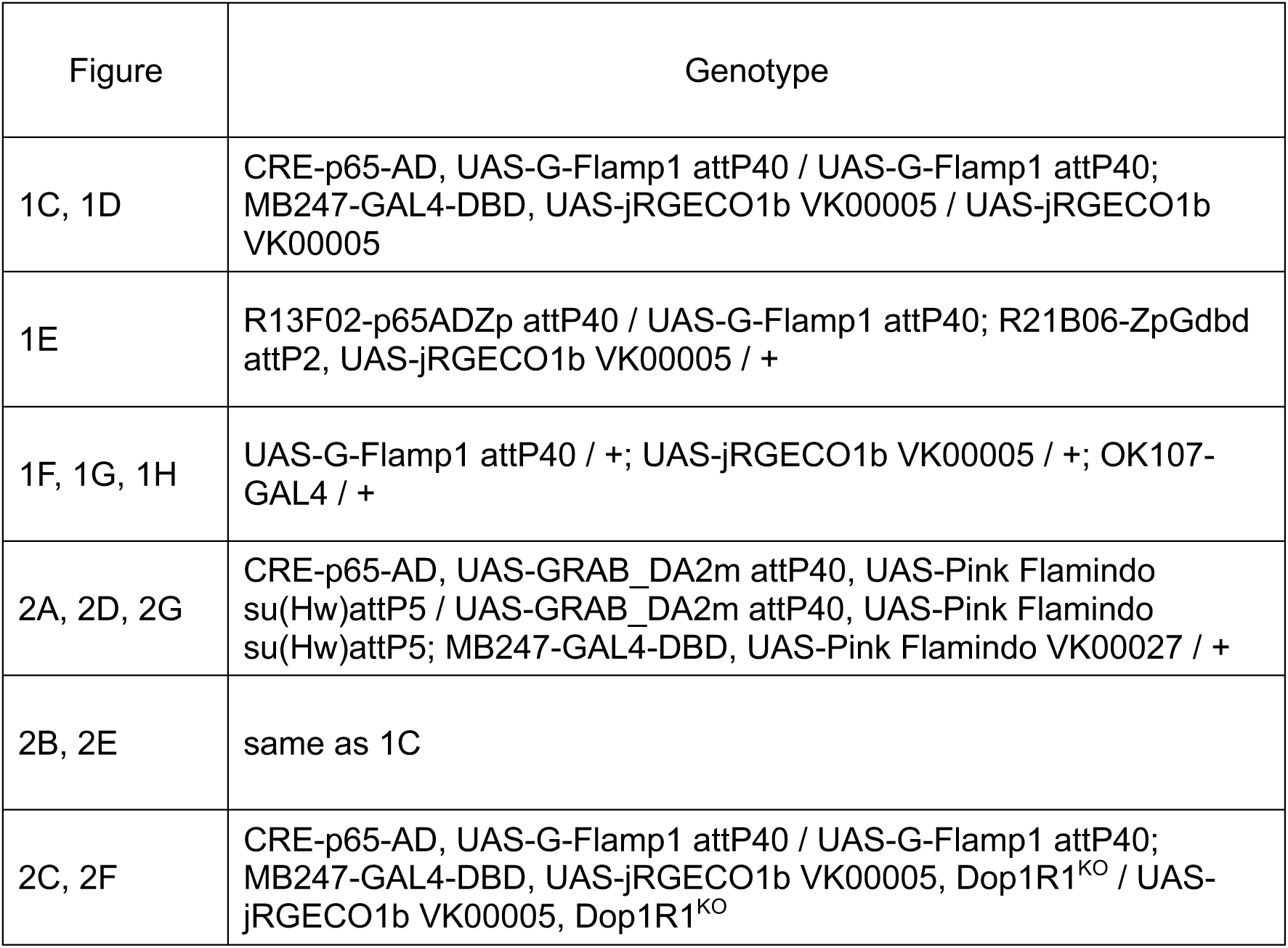
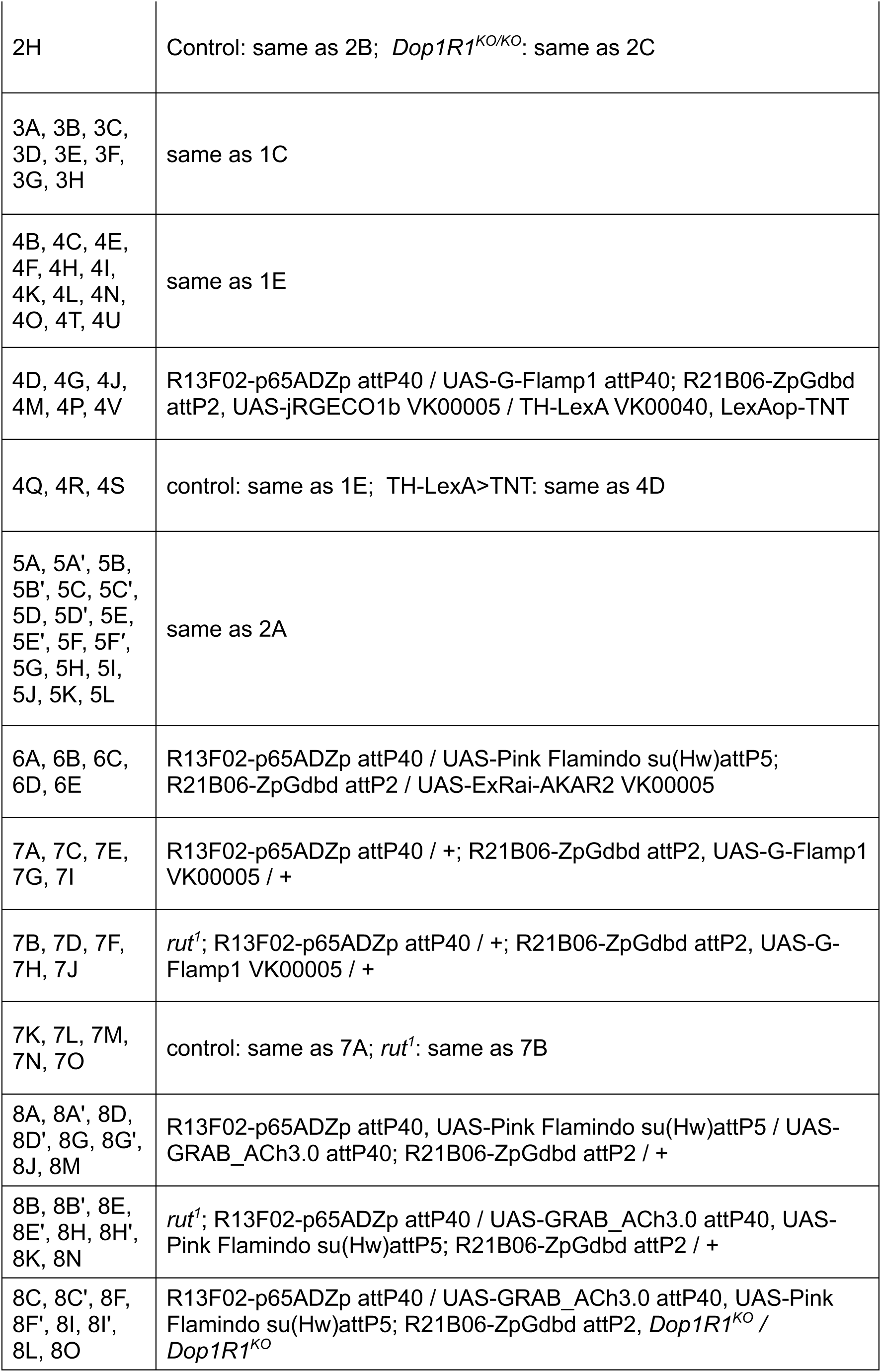
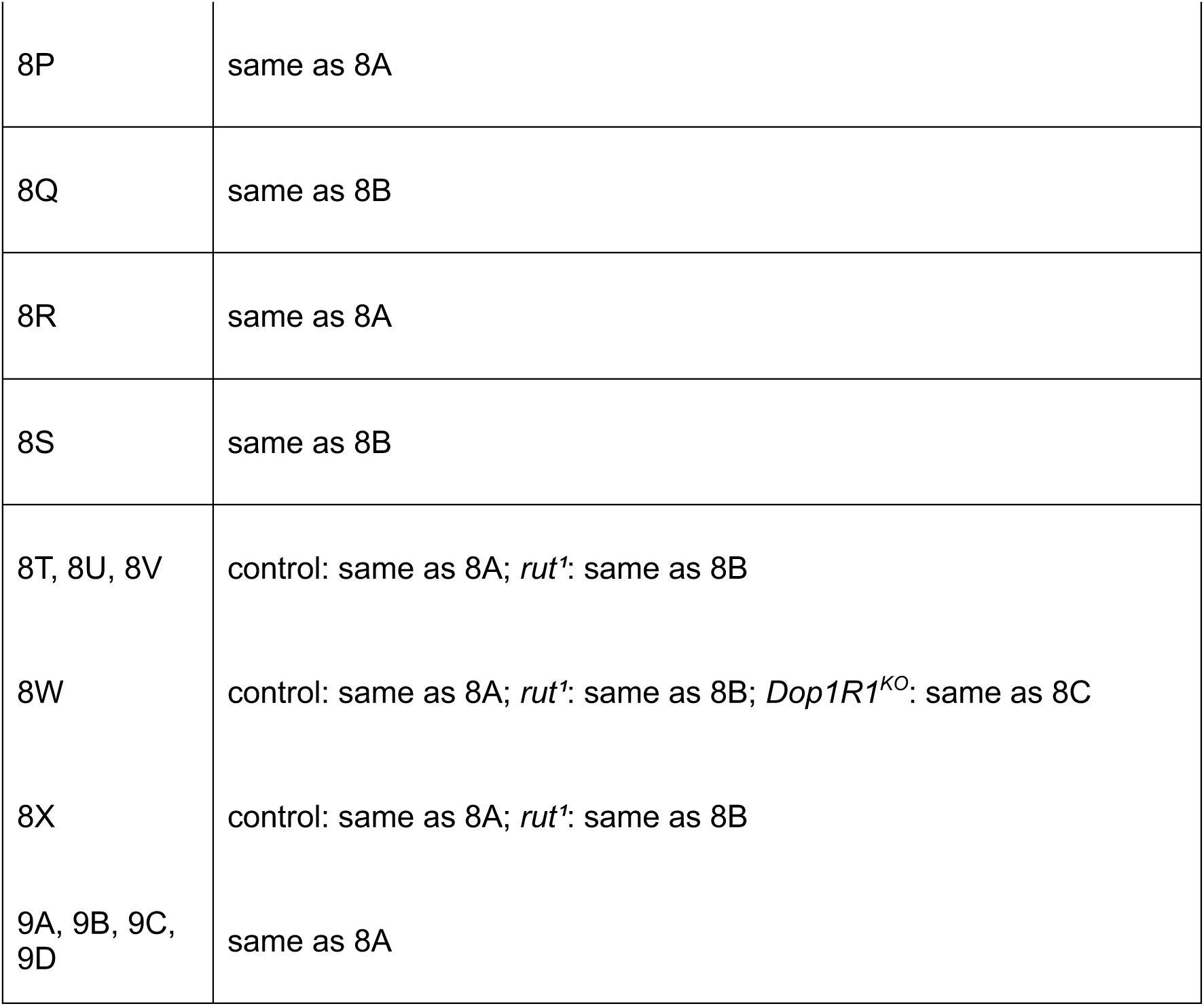

### Construction of UAS-Pink Flamindo and UAS-ExRai-AKAR2

The Pink Flamindo cDNA was amplified by PCR from the plasmid (provided by Kitaguchi, Addgene plasmid #102356). The amplified fragment was then integrated into the 20xUAS vector originating from pJFRC7-20xUAS-IVS-mCD8::GFP (Pfeiffer, Addgene plasmid #26220) using the In-Fusion cloning system (Clontech Laboratories). The ExRai-AKAR2 cDNA sequence was codon-optimized for Drosophila melanogaster (Eurofins Genomics), and the resulting synthesized cDNA was cloned into the 20xUAS vector described above.

### Live imaging

#### Chemicals

The following reagents were used: saline (103 mM NaCl, 3 mM KCl, 10 mM trehalose, 10 mM D-glucose, 26 mM NaHCO3, 1 mM NaH2PO4, 1.5 mM CaCl2·2H2O, 4 mM MgCl2·6H2O, 9 mM sucrose, and 5 mM HEPES; osmolality adjusted to 270–280 mOsm/kg), mineral oil (Fisher Scientific, Fair Lawn, NJ, USA, Cat# O1214), 4-methylcyclohexanol, MCH (Cat# 66360, >98% purity, Fluka, Sigma-Aldrich, St. Louis, MO, USA), 3-octanol, OCT (Cat# 74870, >95% purity, Fluka, Sigma-Aldrich, St. Louis, MO, USA), apple cider vinegar, ACV (Mizkan jun-ringo-su, Mizkan holdings, Japan, ASIN#B000H2SCTW), forskolin (Tokyo Chemical Industry, Cat# F0855), and ethanol (Fujifilm Wako, Japan, Cat# 054-00466).

#### Fly preparation

Unless otherwise specified, female flies aged 5-16 days were used. Flies were mounted on a custom-made imaging dish immediately after brief CO_2_ anesthesia for 3–5 s. The head capsule and dorsal thorax were secured to the dish by light-curing glue (DYMAX Medical Adhesive, 1204-M-SC, USA). The proboscis was also glued onto the capsule to reduce brain movement. A window on the top of the head capsule was opened using sharp forceps (Corte Instruments, USA) and a thin injection needle in saline. Air sacs and fat body tissues were carefully removed, and muscle 16 was cut using forceps to avoid excessive brain movement during imaging.

### Olfactory stimulation

Odors were delivered with a custom-made, 8-channel olfactometer controlled by a custom LabVIEW application (National Instruments). While a fly was mounted under the microscope, a constant stream of pure air (0.4 L/min) was directed toward it. Thirty mL of odor solution was placed in a 100 ml glass bottle (Duran, Cat# TGK 371-05-20-03) with an Omnifit GL45 bottle cap (Diba Omnifit, Cat# OM4212), PTFE tube (Cat# BLS181033, 1/8 in × 2.4mm), and USL-PTFE joint (OOU-095-7×1/8, custom-made, 7mm × 1/8, https://www.u-s-l.co.jp). All odorants except for ACV were diluted 1:10 vol/vol in mineral oil. ACV was diluted 1:10 vol/vol in Milli-Q water. After a stimulation trigger, a solenoid valve redirected a portion of the air stream through the headspace of the glass bottle for 3 s. The solenoid valve was opened from 15 to 18 s after the start of each recording. Because of the transit time through the olfactometer tubing, the odor was estimated to reach the fly approximately 1 s after valve opening, corresponding to an effective odor-delivery period of approximately 16–19 s. Two mass flow controllers (SEC-E40, HORIBA STEC, Japan) were used in parallel to regulate the flow of pure air and odors. The odor stream rejoined the pure air stream 40 cm from the end of the odor delivery tube (4 mm in diameter). The odor was released from a position 5 mm in front of the fly at a flow rate of 0.4 L/min. Subsequently, it was collected through a tube (3 mm in diameter) located 15 mm behind the fly at a flow rate of 2 L/min, using a vacuum pump (ULVAC DA-20D, ULVAC KIKO, Inc.) equipped with a charcoal filter. The intervals between odor exposures were at least 60 seconds.

### Electric shock delivery

Electric stimulation was delivered through two platinum-wire electrodes positioned at the base of the legs and on the abdomen using a micromanipulator (MNM-25, Narishige, Tokyo, Japan). At 17.5 s after the start of recording, a 60-V electric shock was delivered to the fly for 10 ms using an electric stimulator (SEN-3401, Nihon Kohden, Japan) and isolator (SS-203J, Nihon Kohden, Japan). For odor–shock pairing, electric stimulation was delivered 1.5 s after the onset of the 3-s odor presentation. Effective shock delivery was verified using a single 10-ms test pulse outside the conditioning trials by observing leg movements and abdominal contractions with a CCD camera (STC-TB33USB, Omron Sentech, Tokyo, Japan).

### Forskolin bath application

Forskolin was dissolved in a solution containing saline and 100% EtOH in a 3:1 ratio, to prepare a 5 mM stock solution. From this stock solution, a working solution was prepared by dilution with saline to achieve a concentration of 100 μM forskolin in saline with 0.5% EtOH. As a control, saline with 0.5% EtOH was used. For live imaging, the standard saline solution was removed, leaving approximately 50 μL of solution between the objective and the dish.

Then, a syringe containing saline with 0.5% EtOH was connected to the dish, allowing the space between the objective and dish to be filled. The recording was performed for a duration of 4 minutes. Subsequently, the saline with 0.5% EtOH was replaced with 1 mL of the 100 μM forskolin solution in saline with 0.5% EtOH, and another 4-minute recording was conducted. During this time, the responses to odors were recorded. At the end of the recordings, the 100 μM forskolin solution in saline with 0.5% EtOH was washed out by replacing it with the standard saline solution four times, and the responses to odors under the wash-out conditions were also recorded.

### Two-photon laser microscopy

In vivo two-photon imaging was performed using a two-photon microscope (Olympus FVMPE-RS; Olympus, Tokyo, Japan) equipped with a water-immersion objective lens (XLPLN25XWM, 25× N.A. 1.05), a titanium:sapphire pulsed laser (INSIGHT DS DUAL-OL; Spectra-Physics, Santa Clara, CA, USA), and a galvanometric scanner to facilitate live imaging. The emitted light was collected using GaAsP photomultiplier detectors with 495-540 nm and 575-645 nm bandpass filters (FV30-FGR). G-Flamp1 was excited at 925 nm, GRAB_DA2m, GRAB_ACh3.0, and ExRai-AKAR2 were excited at 980 nm, while jRGECO1b and Pink Flamindo were excited at 1040 nm. Using the Olympus FV31S-SW imaging software, we acquired time series images with different dimensions: 400-430 x 120-130 pixels (0.14-0.19 μm/pixel, γ1-γ2 zoom in Figure 3), 275-320 x 140-155 (0.33-0.50 μm/pixel, γ1-γ5 in Figures 1, 2, 4, 5, 6, 7, 8, and 9), and 180-250 x 150-165 (0.15-0.20 μm/pixel, α′3/α3 in Figure 1). The acquisition was performed at a frame rate of either 333 or 500 ms/frame in a sequential line scan mode, ensuring that the time difference between acquiring images for channels 1 and 2 was less than 2 ms. By default, images were acquired for a duration of 40 s to capture the response to each stimulus and were saved for subsequent image processing. The solenoid valve for odor stimulation was activated 15 s after the start of each recording, and delivered the odor for a duration of 3 s, corresponding to frames #46-54 (333 ms/frame) or #31-36 (500 ms/frame). For quantitative analysis, ΔF/F0 images of peak responses were generated by averaging frames #51-65 (333 ms/frame) or frames #35-44 (500 ms/frame). The experimental flies, which were housed in vials with food, were transferred to an imaging room maintained at 22°C at least two days prior to the imaging experiments.

### Image analysis

The image analysis was conducted as described previously (Hiroi et al., 2013), with several modifications. The images within each recording were stabilized using the template matching plugin of Fiji (ImageJ). To mitigate the impacts of detector noise and uncorrected movement, all reporter signals underwent an initial processing step employing a median filter (with a radius of 1 pixel). After stabilization within each recording, images obtained from the same fly were aligned by template matching. Subsequently, ΔF/F0 was calculated using the mean fluorescence intensity during the 1-s interval from 15 to 16 s after the start of the recording as F0. This interval immediately followed valve opening but preceded the estimated arrival of the odor at the fly. A median filter (with a radius of 1 pixel) was applied during this stage. To identify the KC region of interest (ROI), Li’s binarizing algorithm was applied in Fiji to the mean signal intensity over the entire recording. ROIs for each compartment were first manually identified, using the anatomical characteristics as a guide. Because the boundary between the γ3 and γ4 compartments was occasionally difficult to determine, it was pragmatically placed midway between the regions assigned to γ3 and γ4. The drawing and preservation of ROIs, as well as the measurement of ΔF/F0, were performed in a semi-automated manner using both built-in and custom-written macro routines in Fiji. Additional image-processing steps, including pixel-wise correlation analysis and plotting of temporal dynamics, were performed using MATLAB scripts. For analyses of the γ2/γ4 response ratio, datasets in which the γ4 response was negative were excluded because a ratio with a negative denominator cannot be interpreted. For paired comparisons, both measurements from the corresponding fly were excluded if this criterion was met in either condition.

### Scatter plot and correlation analyses

We quantified the pixel-wise relationship between Ca^2+^ and cAMP signals within the γ1 and γ2 compartments using mean stimulus-evoked ΔF/F0 images. For each recording, ΔF/F0 was calculated as described in the preceding Image analysis section. Within each compartmental ROI, the mean ΔF/F0 value was calculated for each pixel over the 1–6 s period following the presentation of OCT alone or OCT paired with electric shock (OCT + shock). Ca^2+^ and cAMP signals were acquired by sequential two-photon excitation in the same optical plane, with an interchannel temporal offset of <2 ms, and were therefore treated as temporally and spatially corresponding at the pixel level. The resulting Ca^2+^ values were plotted against the corresponding cAMP values, with Ca^2+^ on the x-axis and cAMP on the y-axis. Analyses were performed separately for the γ1 and γ2 compartments.

For each sample, Pearson’s correlation coefficient (r) was calculated to quantify the pixel-wise correspondence between Ca^2+^ and cAMP signals. To obtain within-signal self-correlation controls, frames from the same recording were divided into odd- and even-numbered frames. Mean ΔF/F0 images were generated separately from the odd and even frames within the same stimulus-evoked time window, and Pearson’s r was calculated between the corresponding pixel values. The Ca^2+^ and cAMP odd-versus-even self-correlations were used as empirical internal benchmarks for the degree of spatial correspondence detectable under the present imaging conditions.

For comparisons across samples, Pearson’s correlation coefficients were transformed using Fisher’s z transformation. Three paired measures—Ca^2+^ versus cAMP correlation, Ca^2+^ odd-versus-even self-correlation, and cAMP odd-versus-even self-correlation—were compared using the Friedman test followed by Dunn’s multiple-comparisons test. Mean correlation coefficients reported in the Results were obtained by averaging the Fisher’s z-transformed values across samples and then back-transforming the mean values to r. For descriptive purposes, the back-transformed mean r values were also squared. These squared values do not represent the mean R² across samples.

As a complementary analysis, simple linear regression was applied to the same pixel-wise datasets. The slope of the regression line was calculated separately for each sample, and the slopes of the three paired measures were compared using the Friedman test followed by Dunn’s multiple-comparisons test. Comparisons of Fisher’s z-transformed correlation coefficients were regarded as the primary statistical analysis because regression slopes are influenced by differences in signal scaling. Data extraction, scatter-plot generation, Pearson correlation analysis, Fisher’s z transformation, and linear regression analyses were performed using MATLAB. Group-level statistical comparisons were performed using Prism 9.

### Within-sample z-score normalization

For each fly, condition (odor × phase), and signal, the raw values across the four γ-compartments (γ2, γ3, γ4, γ5) in Figures 4, 5, 6 and 8, and the five compartments (γ1, γ2, γ3, γ4, γ5) in Figure 7 were standardized to zero mean and unit population variance within that fly and condition (z-score, ddof = 0): z = (x − μ) / σ, where μ and σ are the mean and population standard deviation of the compartment values. This step removes each fly’s overall signal amplitude and isolates the relative compartmental profile. To visualize how each signal’s compartmental profile evolves over the course of conditioning, the within-sample z-scored values were plotted across the compartments for each phase (Pre, Cond.1, Cond.2, Cond.3, Post), with the mean ± SEM across flies shown at each compartment.

### Pattern correlation

For a given fly, condition, and pair of signals, pattern correlation (r) was computed as the Pearson correlation coefficient between the two signals’ within-sample z-scored four-element vectors. Because both vectors are already standardized to zero mean and unit variance, this is equivalent to r = (1/4)Σ z_1i_z_2i_ (i = 1–4). r = +1 indicates that the two signals place their relative emphasis on identical compartments; r = −1 indicates a mirror-image relationship; r ≈ 0 indicates no relationship between the two compartmental profiles. Individual r values (one per fly per condition) were Fisher z-transformed and compared between Pre and Post using two-sided Wilcoxon signed-rank tests, with each fly serving as its own control. Because responses during Cond.1–3 included the shock-evoked component, these phases were described but not statistically tested. For comparison with shock-evoked profiles (Figure 5K, L), each fly’s OCT-evoked profile was correlated with its own shock-evoked profile for three predefined combinations: DA versus shock-evoked DA, cAMP versus shock-evoked DA, and cAMP versus shock-evoked cAMP. For between-genotype comparisons (Figure 8U, V), the change from Pre to Post (Δz = Post − Pre) and the odor-selectivity contrast (Δz(OCT) − Δz(CS−)) were compared using the Mann–Whitney U test. The within-sample z-scores and pattern correlation coefficients described above were first computed using Python code generated with the assistance of the AI model Claude (Anthropic). The code was then ported to MATLAB, and the values computed in MATLAB were confirmed to agree with those obtained in Python to within rounding error. The resulting values were subjected to statistical analysis using Prism 9 (GraphPad Software), as described in the Statistics section.

### Statistics

Statistical analyses were performed using Prism 9 (GraphPad Software). One mushroom body was analyzed per fly, and n denotes the number of flies. All group-level inferential comparisons were two-tailed and nonparametric unless otherwise specified. The Wilcoxon signed-rank test was used for paired comparisons between two conditions, the Friedman test followed by Dunn’s multiple comparisons test with adjustment for multiple comparisons was used for comparisons involving more than two paired conditions, and the Mann– Whitney U test was used for comparisons between unpaired groups. For the genotype comparisons shown in Figures 7M, 7N, and 8W, responses to MCH, OCT, and ACV were averaged within each fly before statistical analysis. Each fly therefore contributed one value per compartment and sensor, and the fly-level averages were compared using the Mann– Whitney U test (Figure 7M, N) or the Kruskal–Wallis test followed by Dunn’s multiple comparisons test against control (Figure 8W). For unpaired genotype comparisons (Figures 4Q–S, 7M, 7N, and 8W), flies in which only stimulus-evoked responses were recorded without conditioning were also included to increase the sample size. The significance threshold was set at α = 0.05. Details of the statistical tests, exact p values, and the comparisons performed are provided in the figure legends unless otherwise specified. Data in bar graphs are presented as mean ± SEM.

## Supplementary videos

Stacks of ΔF/F0 images corresponding to the averaged peak responses were converted into AVI files using Fiji, assembled in Keynote, and exported in HEVC format.

## Use of AI tools

Claude (Anthropic) was also used for language editing and proofreading of the manuscript; all edits were reviewed by the authors.

## Acknowledgements

We thank Yuuichi Iino, Joshua Johansen, Hokto Kazama, Satoshi Kida, Tom Kornberg, Minoru Saitoe, and Kazumichi Shimizu for their valuable comments. We are grateful to Atsushi Miyajima for providing access to the Ti:sapphire laser. We also thank H. Takishita and M. Sato for excellent help with fly husbandry. We would also like to thank Yoshinori Aso, Chi-Hon Lee, Jun Chu, Yulong Li, Taro Ueno, the Kyoto DGRC, the NIG Stock Center, the Bloomington Stock Center, and the Janelia Research Campus/HHMI for fly stocks. This work was supported by grants from the Ministry of Education, Culture, Sports, Science and Technology (17K17681 and 19K16254 to T.A., 21K06388 to D.Y. and 19H03268 and 24K09543 to T.T.) and the Takeda Science Foundation to T.T.

Author contributions: T.A., D.Y. and T.T. designed the research; T.A. performed all live imaging and statistical analyses; D.Y. constructed transgenes and performed live imaging in the early phase; M.H. developed an imaging data acquisition and analysis system; Y.M. conducted genetic and histological experiments; Y.U. performed statistical analyses. T.A. and T.T. wrote the manuscript.

